# Multi-proxy evidence for the impact of the Storegga Slide Tsunami on the early Holocene landscapes of the southern North Sea

**DOI:** 10.1101/2020.02.24.962605

**Authors:** Vincent Gaffney, Simon Fitch, Martin Bates, Roselyn L. Ware, Tim Kinnaird, Benjamin Gearey, Tom Hill, Richard Telford, Cathy Batt, Ben Stern, John Whittaker, Sarah Davies, Mohammed Ben Sharada, Rosie Everett, Rebecca Cribdon, Logan Kistler, Sam Harris, Kevin Kearney, James Walker, Merle Muru, Derek Hamilton, Matthew Law, Richard Bates, Robin G. Allaby

## Abstract

Doggerland was a land mass occupying an area currently covered by the North Sea until marine inundation took place during the mid-Holocene, ultimately separating the British land mass from the rest of Europe. The Storegga Slide, which triggered a tsunami reflected in sediment deposits in the Northern North Sea, North East coastlines of the British Isles and across the North Atlantic, was a major event during this transgressive phase. The spatial extent of the Storegga tsunami however remains unconfirmed because to date no direct evidence for the event has been recovered from the southern North Sea. We present evidence that Storegga associated deposits occur in the southern North Sea. Palaeo-river systems have been identified using seismic survey in the southwestern North Sea and sedimentary cores extracted to track the Mid Holocene inundation. At the head of one palaeo-river system near the Outer Dowsing Deep, the *Southern River*, we observed an abrupt and catastrophic inundation stratum. Based on lithostratigraphic, macro and microfossils and sedimentary ancient DNA (sedaDNA) evidence, supported by optical stimulation luminescence (OSL) and radiocarbon dating, we conclude these deposits were a result of the Storegga event. Seismic identification of this stratum to adjacent cores indicated diminished traces of the tsunami, largely removed by subsequent erosional processes. Our results demonstrate the catastrophic impact of Storegga within this area of the Southern North Sea, but indicate that these effects were temporary and likely localized and mitigated by the dense woodland and topography of the area. We conclude clear physical remnants of the wave are likely to be restricted to inland basins and incised river valley systems.

---

The Holocene pre-inundation landscape of the southern North Sea, known as Doggerland, was an area associated with Mesolithic hunter-gatherer communities^1^. Sea level rise during the mid-Holocene period at the regional scale was episodic due to local variations in isostatic rebound, autocompaction and palaeotidal range, the precise timing and extent of which, and consequently impact on Mesolithic communities, remains unclear^2^. A key event however during this period was the Storegga Slide, which occurred off the Norwegian Atlantic coast 8.15 thousand years before present (Kya)^2–8^. This has been speculated to have triggered a catastrophic tsunami associated with the final submersion of Doggerland ^2,9^. Despite the apparent magnitude of the Storegga Slide Tsunami evident in northern North Sea, reflected both in sediment deposits and model predictions ^4–7,10^, there has been a surprising lack of physical evidence to suggest the tsunami reached the southern North Sea.^2,9,11^, Figure 1.

**Figure 1.**
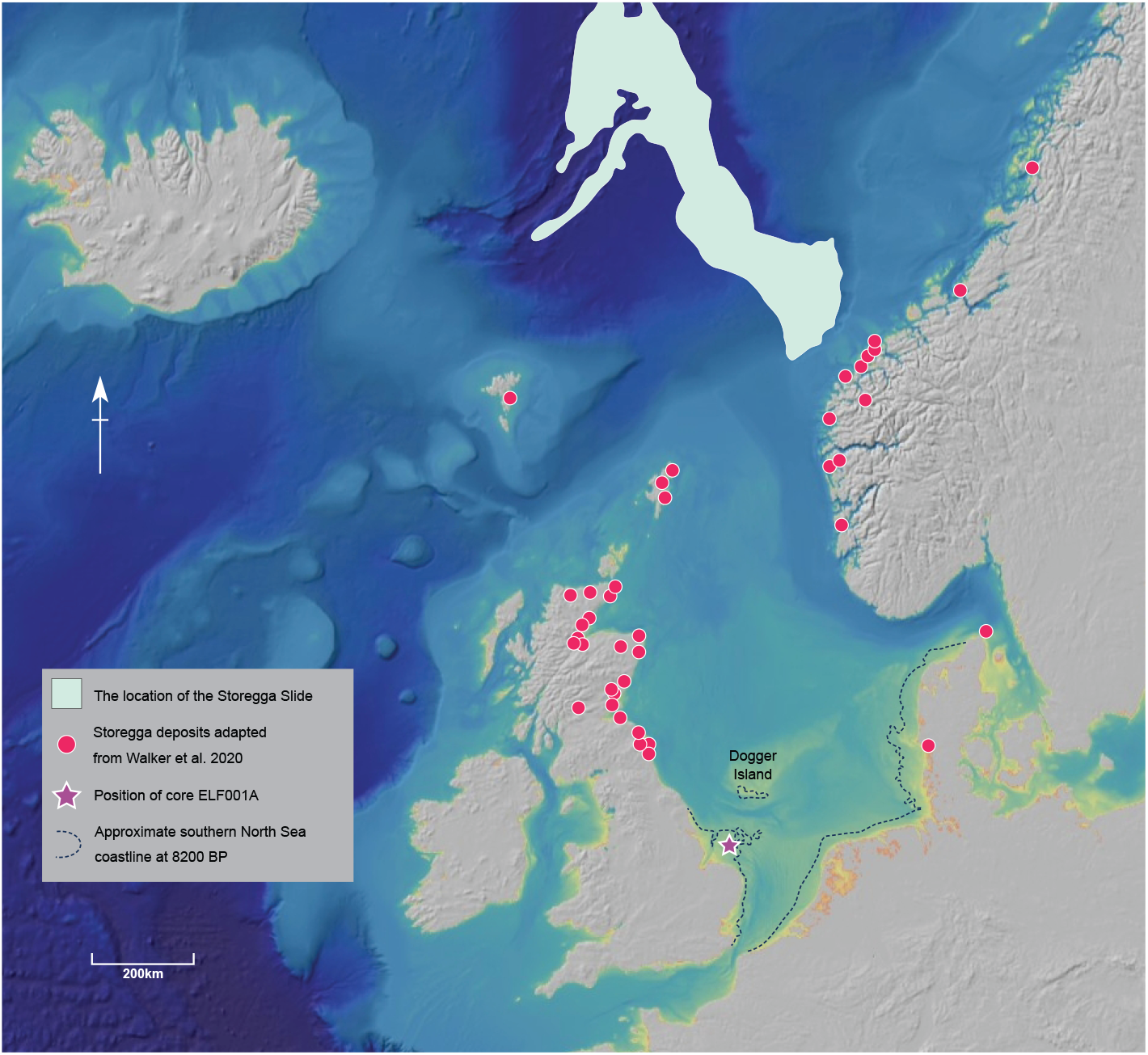
The North Sea, Storegga slips and associated deposits and core ELF001A. Location of deposits associated with Storegga slip follow data in references 6, 9, 10 and summarized in 11.

## Reconstruction of the South Western Doggerland archipelago

In order to reconstruct the early Holocene palaeolandscape of the southern North Sea we used a seismic survey estimating the coastline at 8.2 Kya based on bathymetric contours^12^ using existing sea level curve data^13^ (SI Appendix Text S1). From these data we inferred a generalized landscape, Figure 2. The survey confirmed that the area that had been Doggerland was by this time represented by an archipelago and residual stretches of coastal plain off the eastern coast of England. Within the residual plain were a series of palaeochannels representing river systems within glacial valleys incised through Late Devensian terminal moraines^14^, including a channel associated with the Outer Dowsing Deep, for which we used the term the Southern River. This channel runs north to south terminating in north and south-facing headlands with a central basin. A restricted area around the central basin was associated with a distinct but discontinuous seismic signal suggestive of an anomalously distinct and partially eroded stratum (SI Appendix Text S1.1).

**Figure 2.**
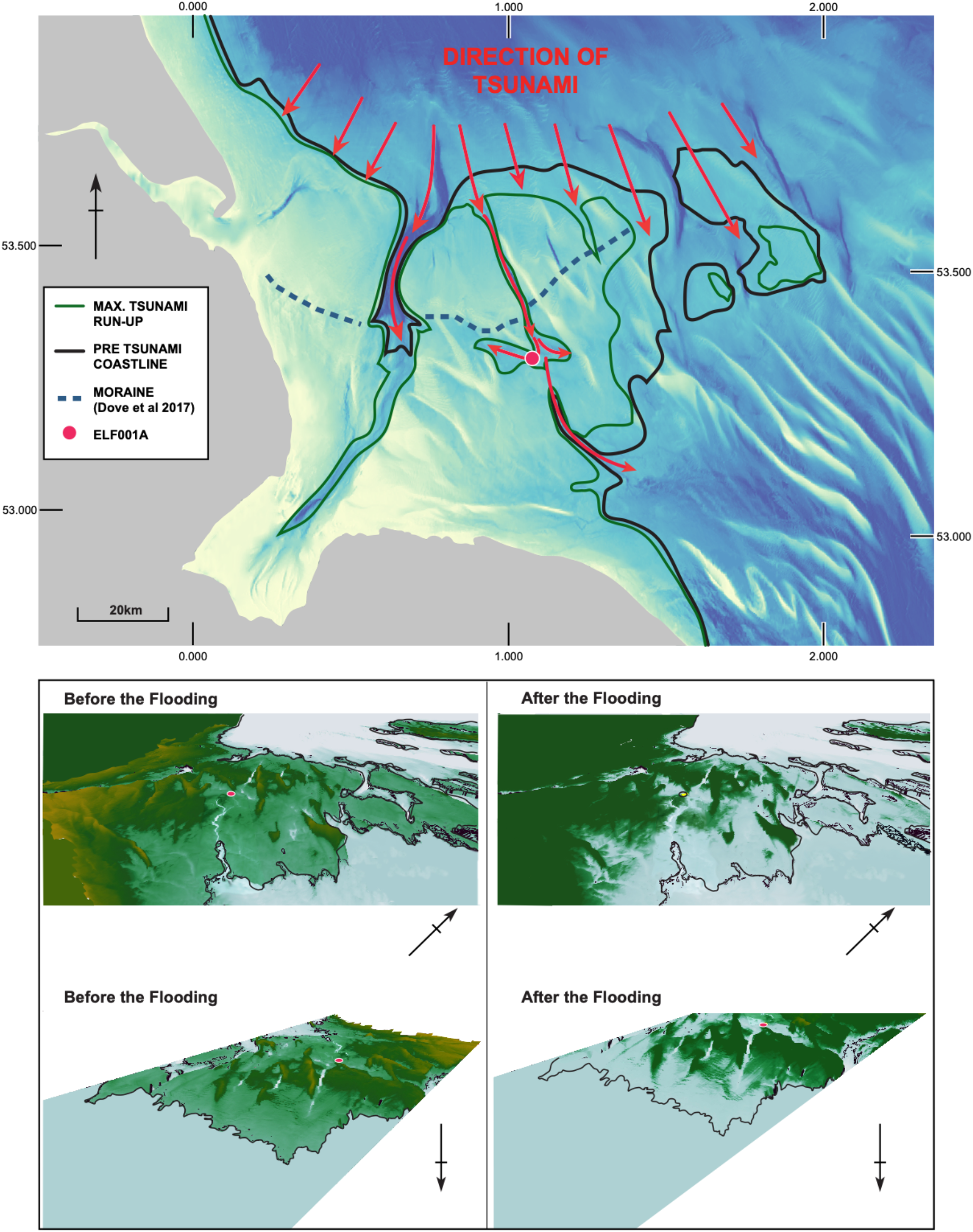
Geormorphological reconstruction of the palaeolandscape off the east coast of England c. 8.2 Kyr BP. Above: Coastline reconstruction at 8.2 Kyr BP and inferred direction of tsunami advance. Inset: Area around coring site of 1A in the Outer Dowsing Deep showing areas unlikely to contain tsunami traces because of high elevations or subsequent erosion. Below: 3D rendering of the area from northern and southern perspectives before and after inundation.

## Identification of a Storegga Slide aged tsunami deposit

Using the palaeobathymetric reconstruction of the landscape as a guide, we investigated the distinct localized seismic signal further and collected 15 vibrocores across the central basin to track the Holocene inundation process. The core most strongly associated with the seismic signal (ELF001A) was characterized by a series of finely laminated silt and clay strata interrupted by a 40cm layer (horizon units 1A-6 and 1A-5) of clastic sediment consisting of stones and broken shells, overlaid by sands (Figure 3, Figure S1.3, Table S1.1). The clastic deposit rests on a sharp, eroded surface in the underlying sediments and exhibits fining upwards within the clay-silt fraction while the coarsest fraction (gravels) occurs in the middle of the unit. Unit 1A-6 is associated with a palaeomagnetic profile which contrasts with the preceding fine silt layer below indicating that it originates from a differing geology (SI Appendix Text S2.1), most likely from outside the local catchment (Figure 3D). An elemental core scan (SI Appendix Text S2.2) confirmed the abrupt change associated with Unit 1A-6 (Figure 3C) with Ca and Sr profiles indicating an influx of marine shells, while Ti and K indicate terrestrial environments towards the base of the core. Similarly, the alkane carbon preference index (CPI), which accounts for the ratio of odd to even carbon chained molecules, indicates a composition dominated by terrestrial forms in the lower core that abruptly switches to an aquatic and marine composition from the event stratum (SI Appendix Text S2.3, Figure 3A). Together, the geochemical and geomorphological evidence indicates that Unit 1A-6 is consistent with a powerful event, but no comparably large stratum was detected in adjacent cores.

**Figure 3.**
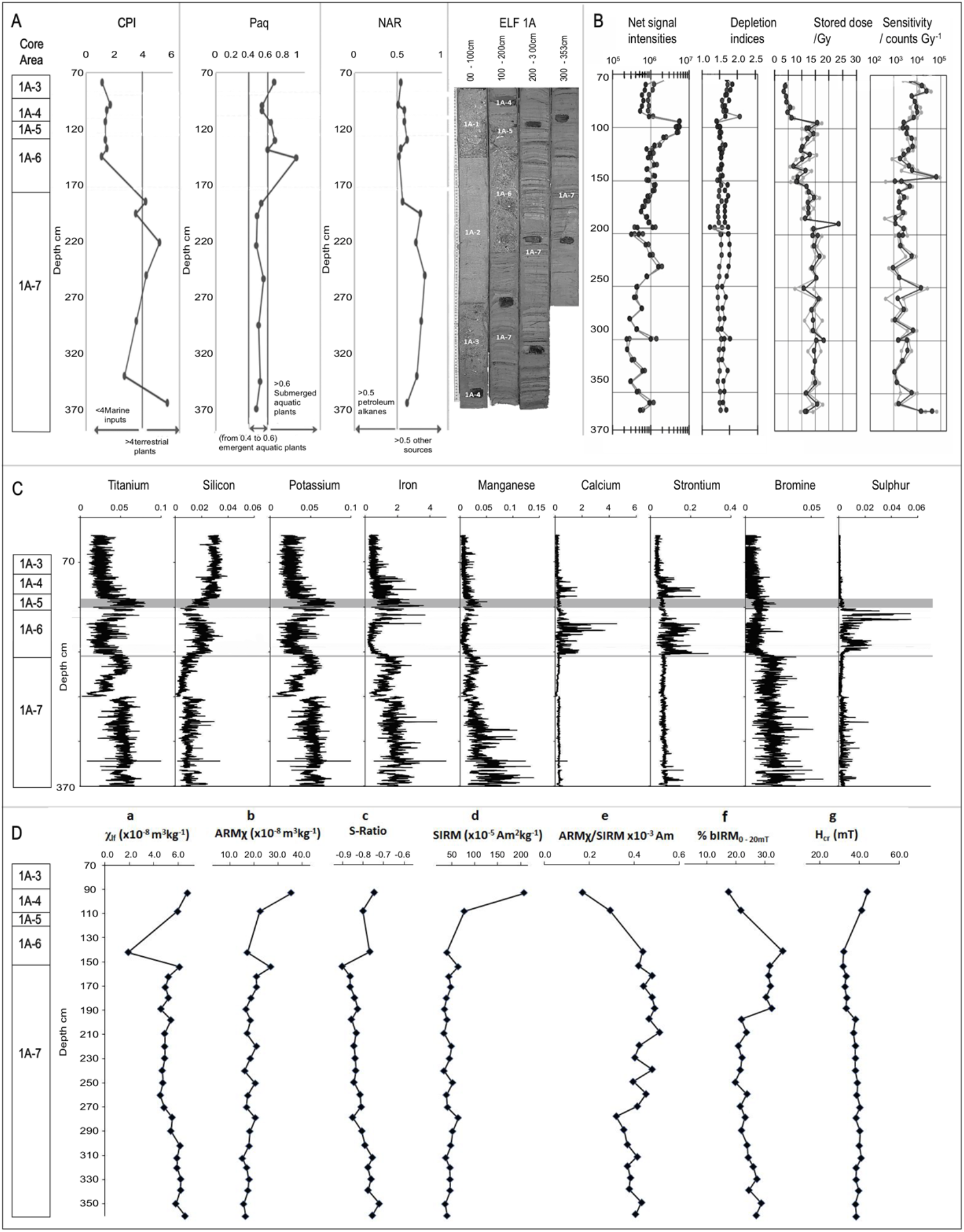
Geochemical analysis of vibrocore *ELF001A*. A: Organic chemistry profile. Carbon Preference Index (CPI) is an indicator of marine or terrestrial environments. N-alkane proxy ration (Paq) distinguishes emergent and submerged aquatic plants. Natural alkanes ratio (NAR) distinguishes n-alkanes from natural oil sources from other sources such as higher plants. B. Optical stimulation lumiscence (OSL). C. Elemental core scan. Horizons associated with marked events are denoted by thin and thick grey lines respectively. Thin line: sudden influx of calcium, strontium and sulphur coinciding with the onset of tsunami associated unit 1A-6. Thick line: Titanium and potassium increases indicate a return to terrestrial influence. D. Palaeomagnetism profile

We investigated the age and depositional rate of the sequence in *ELF001A* using optical stimulation luminescence (OSL), Figure 3B (SI Appendix Text S3.1), and directed AMS radiocarbon dating (SI Appendix Text S3.2). The luminescence profile shows a steady accumulation of signal over time either side of the clastic stratum reflecting a stable sedimentation regime (see Supplementary Information), but shows a disturbed inversion within the stratum consistent with an influx of extraneous matter, Figure 3B. OSL dates could be retrieved under steady sedimentation rate conditions indicating the tsunami deposit occurred at 8.14 ± 0.29 Kyrs. BP (SI Appendix Table S3.1), making this stratum closely contemporaneous to the Storegga Slide. Given the OSL profile, the extraneous matter in this stratum likely reflects material of various ages dredged up from older strata in the surrounding area. This was confirmed by radiocarbon dates from shell fragments within the clastic stratum dated to 8.34 ± 0.3 Kyrs BP (SI Appendix Text 3.2), two centuries prior to the Storegga Slide Tsunami. Together, this evidence convincingly establishes the clastic stratum in core ELF001A as a deposit resulting from the Storegga Slide Tsunami event.

## Palaeoenvironmental impact of the Storegga Slide Tsunami

We further confirmed the catastrophic nature and source of this stratum using multiple palaeoenvironmental proxies, Figure 4. Proxies included foraminifera, ostracods (SI Appendix Text S4.1), pollen (SI Appendix Text S4.2), diatoms (SI Appendix Text S4.3), molluscs (SI Appendix Text S4.4) and sedimentary ancient DNA (sedaDNA), (SI Appendix Text S4.5). Relatively low levels of cytosine deamination and fragmentation patterns consistent with ancient DNA of this age and environment^15^ were observed firstly by mapping sedaDNA to *Quercus*, *Corylus* and *Betula* genomes applying conventional mismatching approaches^16^ (figures S4.2 and S4.3), and secondly applying a novel metagenomic assessment methodology in which all sedaDNA is assessed for deamination damage, which may be more suitable for this data type (Figure S4.4). We then further tested sedaDNA for stratigraphic integrity to assess possible biomolecule vertical movement in the core column (SI Appendix Text S4.5). Figure 4 shows that the sedaDNA demonstrates highly significant differentiation between strata indicating a lack of movement post deposition. Together, these tests indicate that authentic sedaDNA was retrieved and most likely represent the original depositional environment. Interestingly, the same stratigraphic tests applied to pollen generally show a lack of differentiation between strata indicating both a consistent influx of pollen from the surrounding area from oak, hazel woodland, and that the sedaDNA derived from sources other than pollen as has been previously suggested in other sedimentary contexts^17^. This suggests a taphonomy in which the sedaDNA represents a local signal relative to a more regional palynological signal.

**Figure 4.**
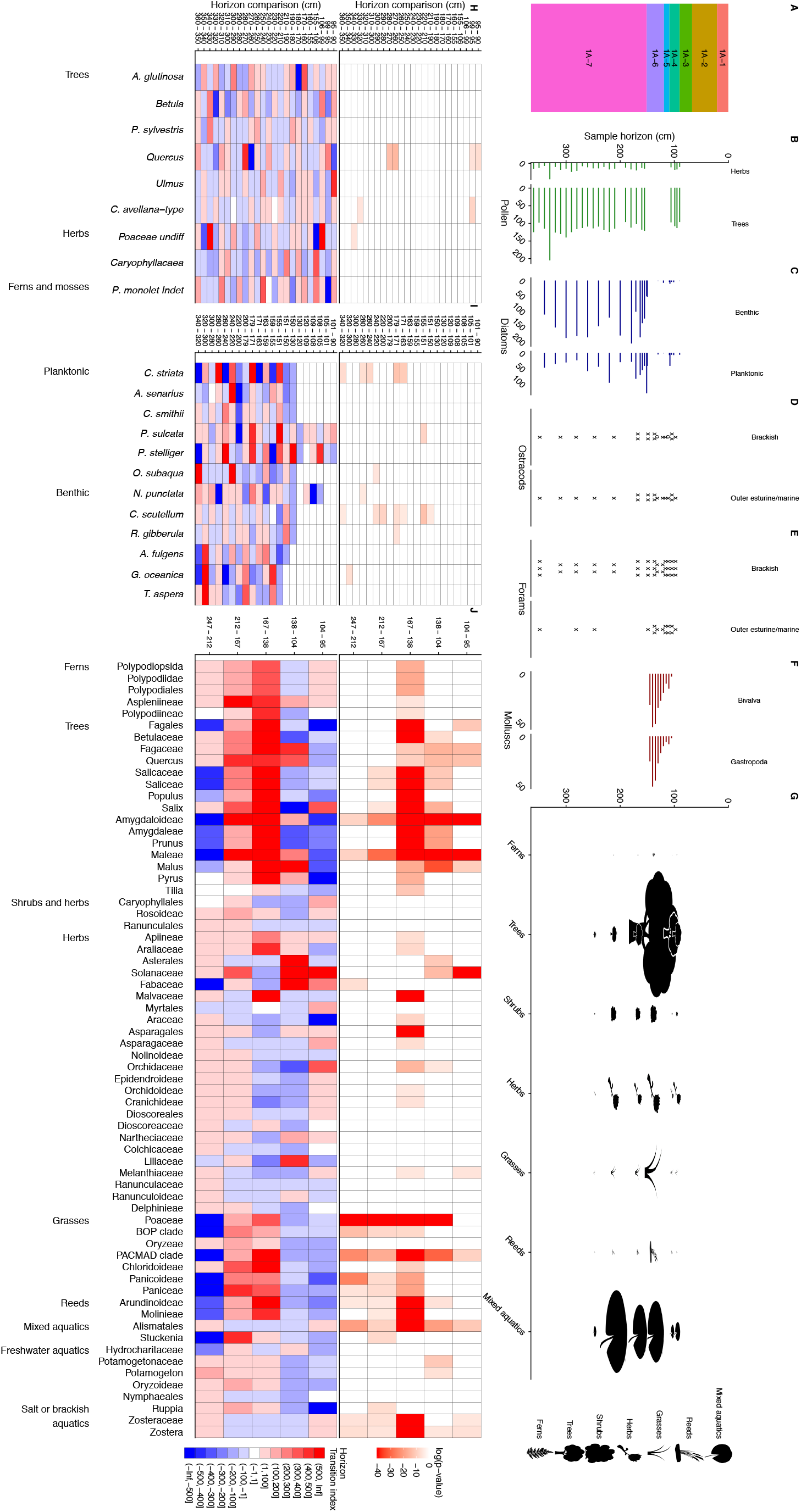
Palaeoenvironmental reconstruction of *ELF001A*. All measurements were taken from lithological units 1A-4 and below. A. Lithological units based on sediment types described in Supplementary Fig 1 and Supplementary Table 1. Panels B-G palaeoenvironmental proxy abundances, for detailed taxonomic break downs see Supplementary Fig. 10 for panels B, C and F, Supplementary Fig. 11 for panels D and E, and Supplementary Fig. 12 for panel G. B. Pollen counts. C. Diatom counts. D. Ostracod abundance E. Foramifera abundance. Foraminifera and ostracods are recorded: o – one specimen; x - several specimens; xx – common; xxx – abundant. F. Mollusc counts. G. Plant sedaDNA vegetation type with icon size relative to biogenomic mass. Panels H-K Assessment of taxon change between sample horizons of taxa with abundances > 50, see Supplementary Information for methods. Below: Index of change between horizons based on changes in maximum likelihood estimators of the probability taxon being selected from each horizon. Blue indicates a decrease in probability moving up the core, red an increase. Above: Probability of observed taxa counts between pairs of horizons being drawn from the same distribution. H. pollen data. I. diatom data. J. sedaDNA data.

Unit 1A-6 is characterized by an abrupt change in both microfossil and sedaDNA evidence. There is an absence of diatoms and pollen; an increase in outer estuarine or marine taxa of ostracods and foraminifera; the appearance of fractured molluscan shells from different and incompatible habitats including sublittoral, intertidal and brackish species; and the sudden and significant influx of all woody taxa in the sedaDNA profile (figures S4.1, S4.5 and Table S4.1). A novel measure of relative biomass, biogenomic mass, based on sedaDNA and genome size (SI Appendix Text S4.5), suggests a higher biomass of trees than either *Zostera* or *Potamogeton* in this stratum, although these latter taxa dominate in other strata, Figure 4 (Figure S4.5). Together, these proxies indicate a violent event that brought with it the debris of surrounding woodland, consistent with the impact, and backwash, of a tsunami that dates to that of the Storegga Slide.

The environment prior to this dramatic event and recorded in the underlying stratum Unit 1A-7 was an estuarine marsh typified by predominantly benthic epiphytic and epipelic diatom communities and brackish foraminifera and ostracods, with a sedaDNA floral profile of *Zostera* and *Potamogeton* as well as members of the Hydrocharitaceae and Araceae present. A small meadow influence is also apparent in the sedaDNA profile including buttercups, orchids, mallows and asterids suggesting proximal open terrestrial systems, Figure S4.5. After the tsunami in units 1A-4 to 1A-1 the foraminifera and ostracod signal indicate a return to estuarine mudflats with a greater abundance of marine taxa such as *Ammonium batavus* indicating a more established marine signal than prior to the event, Table S4.1. The sedaDNA signal also indicates estuarine taxa such as *Zostera*, and a meadow influence, although the biogenomic mass appears greatly reduced suggesting more distant proximity of the flora. A faunal signal considerably weaker than the floral was present throughout the core, but shows a significant elevation in count towards the top units (*p* = 1.0014E-06), indicating the presence of rodents and larger animals such as bear, boar and cloven hoofed ruminants, as well as higher orders of fish (Acanthomorpha, Eupercaria, Osteoglossocephalai), Figure S4.6.

These data lead us to conclude the effects of the tsunami altered the immediate landscape, perhaps opening up the surrounding forest to larger animals, but given the return of the terrestrial signal in subsequent strata we conclude that the final marine inundation occurred at a later time in this area.

## Geomorphological influence on the tsunami propagation

The occurrence of a Storegga tsunami deposit at *ELF001A* 42 km from its contemporary coastline (Figure 2) is unexpected given previous models have estimated a magnitude of wave that would reach 21 km inland in this area^18^ before becoming impeded by a glacial moraine belt^14^. Our reconstruction at 8.2 Kya suggests that the geomorphology of the landscape was likely instrumental in propagating the wave inland. The orientation of the coastline relative to the direction of travel is consistent with a funneling of the tsunami into glacial tunnel valleys that breach the moraine, of which the Outer Dowsing Deep is one, Figure 2. The passage itself is a steep sided U-shaped valley that becomes progressively deeper southwards towards the central basin, which being below the 8.2 Kya sea level supports the palaeoenvironmental proxy evidence for an estuarine system. Such a configuration is expected to lead to an intensification of the wave and consequently a significant impact^19^.

To explore this scenario further we tracked the probable height of the tsunami stratum from *ELF001A* across the landscape using Glacial Isostatic Adjustment models^13^, seabed mapping and Geographic Information System analysis (SI Appendix Text S1.1 and S1.3) and then used seismic data in order to predict where similar deposits would be expected to be preserved in other cores, Figure S1.1. The analysis indicated an absence of strata contemporaneous to the tsunami layer in many areas reflecting widespread erosion subsequent to the tsunami event, Figure S1.2. However, the seismic signal associated with the change in sediment density that was identified as the tsunami stratum in *ELF001A* was used to trace other core locations likely to contain signals originating from the same source, Figure S1.2. Using this technique two other tsunami candidate cores were identified, *ELF003* and *ELF059A*, situated within the central basin area and southern river channel, figures S1.1 and S1.3. It should be noted that this signal was not replicated in other seismic surveys, and the signal, which correlates in this instance cannot, as such, be regarded as diagnostic for the purposes of identifying the presence or absence of similar deposits elsewhere. The identification of the Southern River system by palaeobathymetry allowed the identification for two further cores *ELF031A* and *ELF039*, as containing these deposits in the likely outflowing channel to the south of the basin (SI Appendix Text S1.3). Significant surges in woody taxa similar to that seen in Unit 1A-6 at the predicted tsunami height were detected in *ELF003* and *ELF0039* based on sedaDNA. The corresponding height in *ELF0059A* occurred at the base of the core but was also associated with a significantly higher abundance of woody taxa than overlying strata, and *ELF0031A* generally yielded too little sedaDNA for interpretation, Figure S4.8. Radiocarbon dates from *ELF003* confirmed the corresponding tsunami height to be contemporaneous with Unit 1A-6, supporting the notion that these cores carry a much-diminished signal of the tsunami event.

## The Storegga Slide Tsunami in the southern North Sea and final inundation

Together, these data lead us to conclude that the current physical evidence for the Storegga tsunami in the study area is highly localized because of the channeling effect of the tunnel valley systems and barriers to wave propagation provided by the wooded glacial moraines. This restricted distribution was further reduced by later erosive processes. It may be the case that physical evidence for the Storegga tsunami in the southern North Sea has not been previously observed because it only resides in restricted locales, such as incised river systems, where the geography and local conditions were favorable for preservation.

Our evidence shows that the Storegga Slide Tsunami impacted coastlines in the area of the southern North Sea covered by this study. In coastal areas where human populations may have resided for most of the year, settlement would have been badly affected. The multiproxy evidence suggests the landscape recovered temporarily and hence confirms that the final submergence of the remnant parts of Doggerland occurred some time after the Storegga Slide Tsunami. At the same time, the remaining local terrestrial landscape is suggested to have been more open. Occupation could therefore have continued after the tsunami retreated, but within a much modified coastal landscape before early-mid Holocene eustatic sea-level rise was responsible for finally submerging the remnant Doggerland lowlands and its associated Mesolithic communities.

## Methods

All methods are described in the supplementary information texts. Geomorphological analysis including seismics, coring and palaeobathymetry are presented in SI Appendix S1. Geochemistry analysis including palaeomagnetics, elemental core scans and organic chemistry profiling are presented in SI Appendix S2. Dating of sediments using OSL and organic materials using radiocarbon is presented in SI Appendix S3. Palaeoenvironmental analysis including foraminifera, ostracods, pollen, diatoms, molluscs and sedaDNA are presented in SI Appendix S4.

## Acknowledgements

This project has received funding from the European Research Council (ERC) under the European Union’s Horizon 2020 research and innovation programme (ERC funded project No. 670518 LOST FRONTIER, https://europa.eu/european-union/index_en, https://lostfrontiers.teamapp.com/). The project gratefully acknowledges the support of and the Estonian Research Council (https://www.etag.ee/en/estonian-research-council/ Grant number: PUTJD829) and PGS (https://www.pgs.com/) through provision of data used in this paper under license CA-BRAD-001-2017.

## Supplementary Materials

### Supplementary Text S1 Geomorphology analysis

#### S.1. Seismic survey

The high resolution seismic geophysical dataset was acquired between October 2008 and March 2009 as two separate surveys by Gardline Surveys Limited. The data was obtained by the Gardline Vessel *Vigilant*, which was equipped with a surface-towed boomer system consisting of an Applied Acoustics 300 Plate powered by an Applied Acoustics CSP 1500 Pulse Generator. The receiver consisted of a 12-element single channel hydrophone eel recorded with a Gardline 2012. Digital data logging and initial processing was accomplished using an Octopus 760 geophysical acquisition package (CodaOctopus). During acquisition a swell filter was applied to the data when necessary to correct for the effects of sea swell. The system was operated at a power level of 300 joules with a 350-millisecond fire rate. This equipment setup was used on all profiles with useful data generally recovered to a depth in excess of 25 metres below seabed. The data was initially inspected, and processing accomplished using both SonarMap (Chesapeake Ltd) and GeoSurvey (Coda Ltd) with further processing utilising IHS Kingdom. A number of post-acquisition processing steps were applied to the data which included bandpass filtering, time varied gain and running-sum amplitude gain correction. Sub-surface layers were first-break picked from refracted seismic signals where evident above background noise.

The seismic reflection later associated with the Tsunami deposit was characterized by a negative amplitude of response from -18,000 to -26,000 as well as a sharp phase transition within the wavelet of response from -165 to +171 degrees. This signal was a distinct response within the seismic line and occurred broadly along the same time interval, between 0.035 and 0.037 seconds, indicating a distinct stratum. The signal was not repeated in any of the deeper parts of the seismic line, nor observed in seismic lines outside of the basin surrounding core ELF001A (Figure S1.1). This signal was therefore deemed to be a unique character to this area of the survey. Because of the unique nature of the signal, coring this unit (ELF001A) was undertaken in 2016 to determine the origin of this reflection (Figure S1.2), whereupon subsequent lithological, environmental proxy and dating analyses established the presence of a tsunami deposit (see following methods). Once the reflection had been identified as a tsunami deposit within the core, the seismic signal was re-examined and the correlation between reflector depth against the deposit depth was verified.

When the seismic line was examined, similar responses were noted in patches within three distinct geographic areas within the basin, one 1.96km along the line to the East, a central section 1.40km along the line, where ELF001A was recovered, and a smaller western section 540m along the line (Figure S1.2A.). After the recovery of ELF001A, the discontinuous nature of this reflector was interpreted to relate to post tsunami erosion, occurring during the inundation and submergence of this area. Using this knowledge, cores ELF003 and ELF0059 were identified as candidates to also include traces of the tsunami deposit (Figure S1.1). These patches of similar seismic response were recorded during the re-examination of the data to assist interpretation and guide future survey in these areas.

**Figure S1.1.**
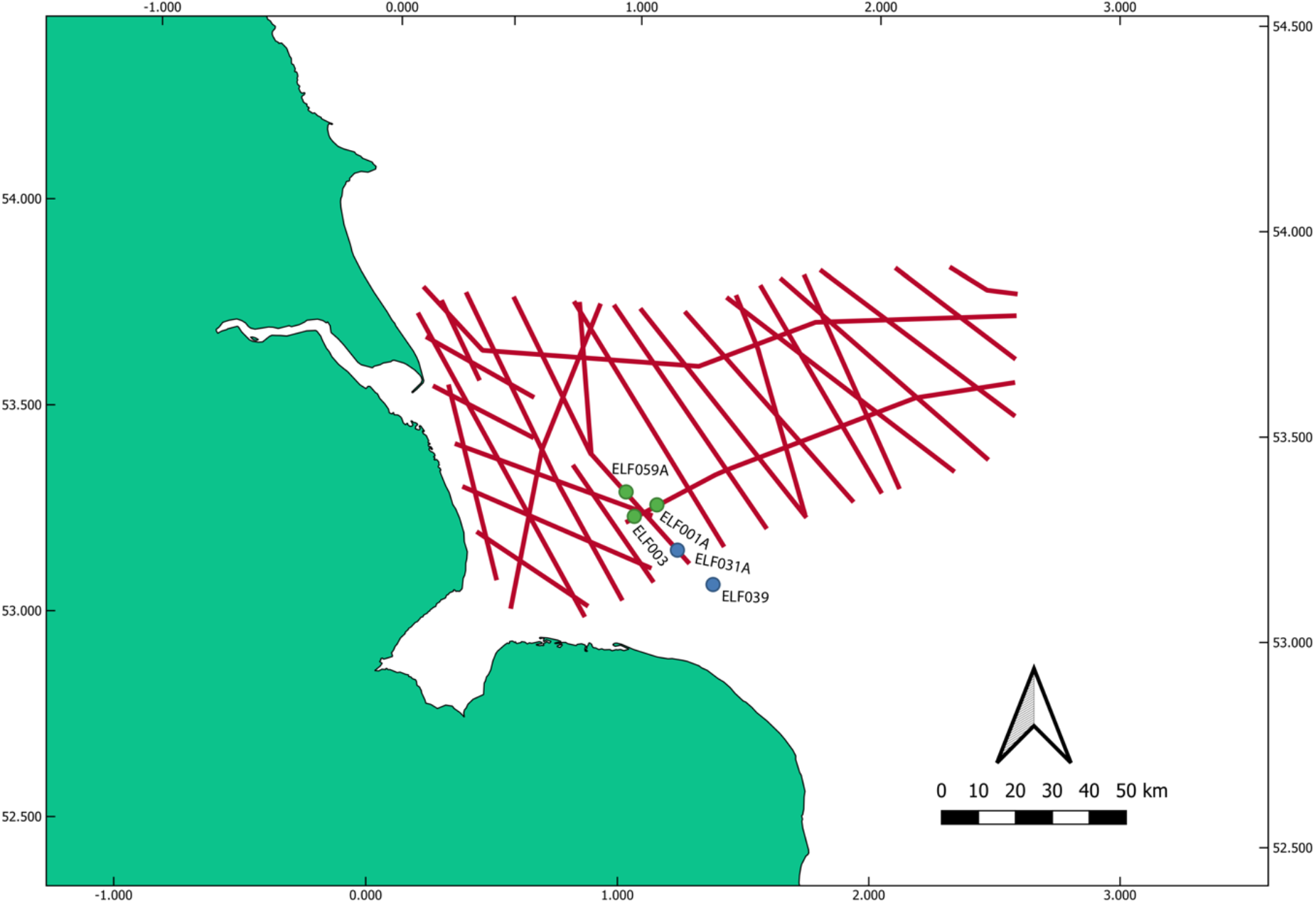
Location of the 2D seismic survey lines and tsunami associated core locations. Cores identified with Tsunami material within the basin are marked green, whilst those identified to be associated with the drainage via the southern river are marked blue

**Figure S1.2.**
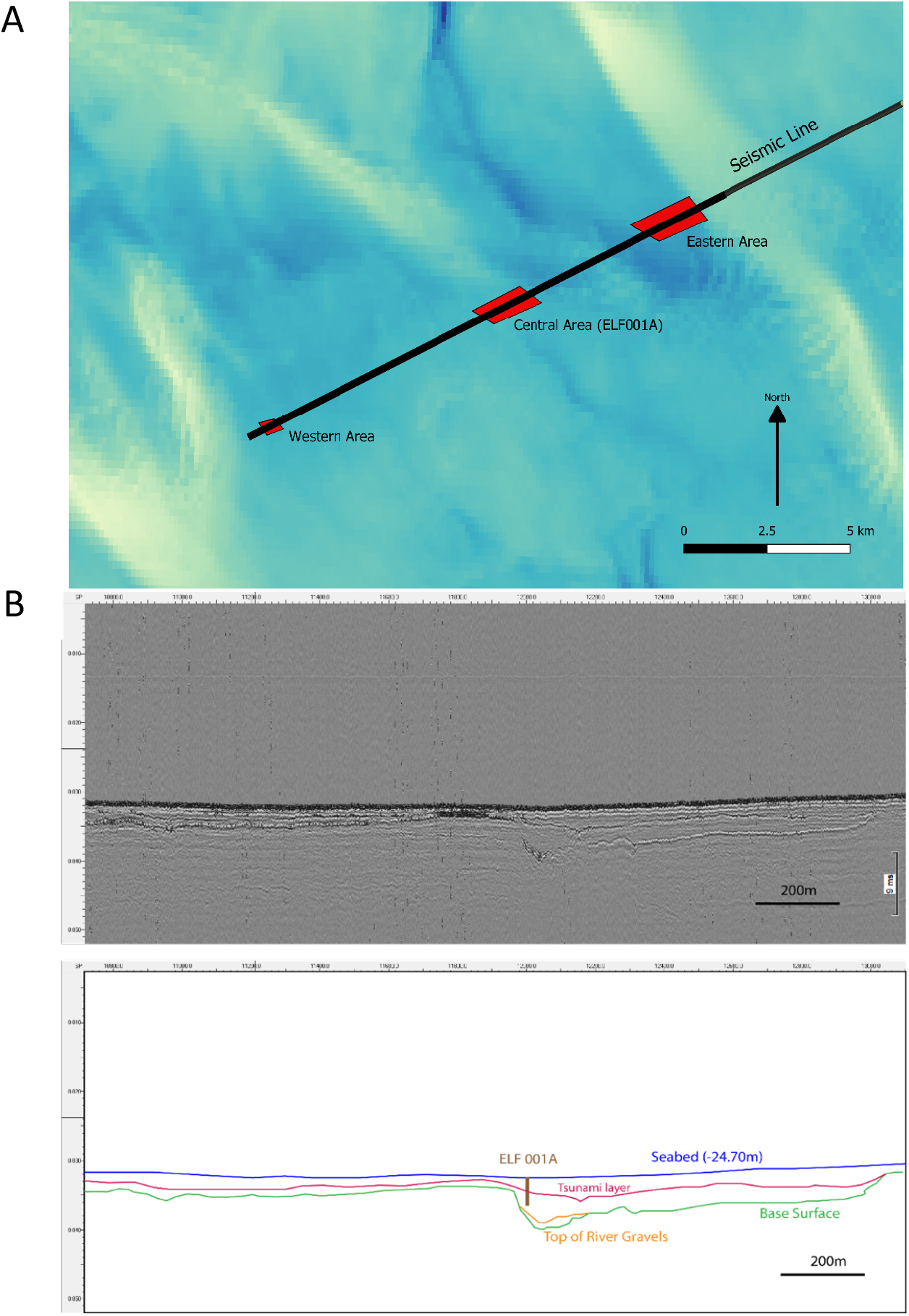
Localized discontinuous seismic tsunami signal. (A). Geographical location of seismic responses (shown in red) along the seismic survey line (shown in black) which show a similar seismic response to that shown by the Tsunami deposit at ELF001A. (B) Upper panel: 2D seismic profile of the central area over the ELF001A location. Lower panel: annotated interpretation of seismic data in upper panel.

#### S1.2 Core acquisition and lithographic assessment

The coring was undertaken as a dedicated survey in September 2016 by Gardline Surveys Limited. A Gardline Geosciences 5 m vibrocorer was used to collect continuous 86 mm diameter samples from 20 sites within the survey area. Opaque liners were used in all cases as optically stimulated luminescence (OSL) dating was required. The cores were sealed and wrapped in black plastic immediately upon recovery. Of the 20 sites sampled 6 required a 2nd attempt to acquire an acceptable sample. Core sites were located upon existing seismic lines to facilitate core correlation with the data. The coring locations were determined following interpretation by the archaeological team and sited over areas of archaeological interest and/or locations with potential for good archaeo-environmental preservation.

Core cutting was carried out under controlled conditions in University of Warwick laboratories. Cutting of cores and initial sampling for preserved sedimentary DNA (sedaDNA) was undertaken in environmentally controlled conditions and under red light so as to minimise likely light contamination of sediments designated for Optically Stimulated Luminescence (OSL) profiling and dating. After cutting, one half of the core was immediately sealed in black plastic for OSL dating. The other half of the core was rapidly assessed, and samples taken for specialist analyses. Core recording procedures follow the guidelines of Jones *et al*. (1999)^20^. The Basic lithostratigraphic profiles from the core logging were drawn as sections to facilitate interpretation and presented via the standard geological modelling software Rockworks (https://www.rockware.com/product/rockworks/).

The initial lithological assessment of core ELF001A revealed seven different sediment types (units) separated by sharp, abrupt or diffuse contact, Table S1, Figure S1.3. Unit ELF001A-6, characterized by structureless, loose medium sands including stones and broken shells consistent with a storm surge, was later established to be consistent with a Storegga Slide Tsunami deposit.

**Figure S1.3.**
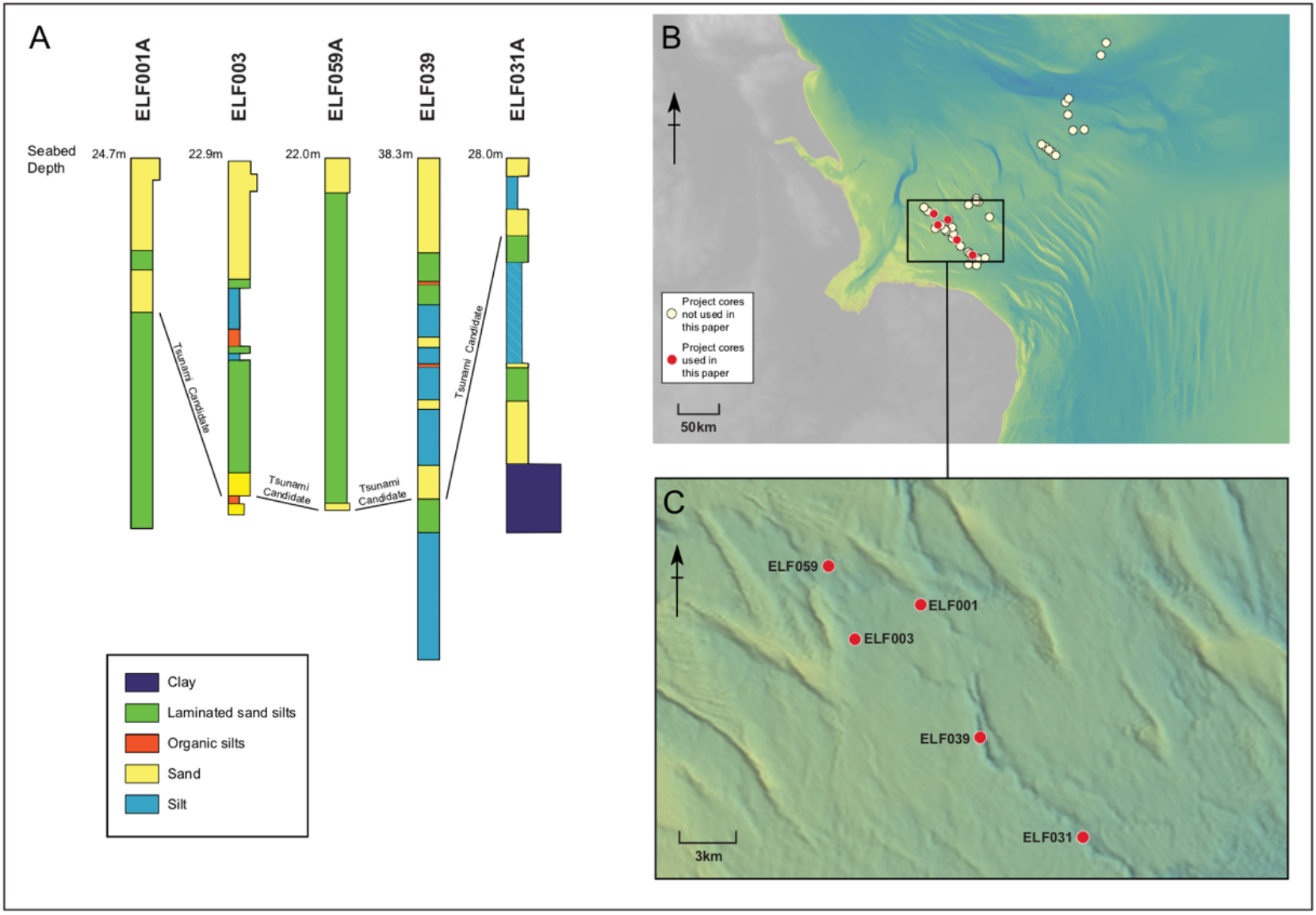
Lithography of candidate Tsunami associated cores. A. Lithographic profiles of cores ELF001A, ELF003, ELF0031A, ELF0039, ELF0059A. B. Total sites cored in study. B. Core locations used in this study.

**Table S1.1.** Lithological profile of sediment core ELF001A.

| Depth below ground surface (m) | Depth O.D. | Lithological description | Unit Number |
| --- | --- | --- | --- |
| 0.00 – 0.21 |  | Yellow shell gravel with whole and fragmented shell. Small sub-rounded to sub-angular gravel clasts (<1cm). Structureless and loose/unconsolidated.<br>---Sharp Contact--- | 1A-1 |
| 0.21 – 0.67 |  | Pale greyish yellow medium sand. Structureless. Occasional small (2-4mm) shell fragments. Some black flecks below 0.46m.<br>---Abrupt Contact--- | 1A-2 |
| 0.67 – 0.90 |  | Greyish yellow medium sand with common shell fragments. Some possibly articulated Mytilus shells. Some burrowing evident, possibly very crudely bedded. Loose and unconsolidated. Some small stones.<br>---Abrupt Contact--- | 1A-3 |
| 0.90 – 1.09 |  | Mid to dark grey very well laminated fine sands and silts. Predominantly sub-horizontal laminations of 2-4mm thick. Occasional dipping laminations of sand. Occasional shell fragments. Moderately cohesive.<br>---Abrupt Contact--- | 1A-4 |
| 1.09 – 1.19 |  | Dark grey silty fine sand with common shell fragments. Shells are <4mm commonly but with occasional fragments larger than 3cm. Shells are broken and sharp/fresh. Possibly some crude bedding. Moderately firm and compact.<br>---Diffuse Contact--- | 1A-5 |
| 1.19 – 1.51 |  | Grey medium sand with very common shells fragments including whole shells and freshly broken shell fragments. Small stones throughout unit. Loose, unconsolidated and structureless.<br>---Sharp Contact--- | 1A-6 |
| 1.51 – 3.50 |  | Mid to dark grey finely laminated silts and fine sands. Sub-horizontal laminations from 2-3mm thick to 1cm thick. Occasional brown organic fragments. Moderately firm and compact. Occasional sand beds 2cm thick. Possibly becoming more silty with depth.<br>---Base 3.50m--- | 1A-7 |

#### S1.3 Palaeobathymetry and estimation of 8.2Kyr shoreline and tsunami run-up

To understand the nature of this Tsunami deposit further, the seabed bathymetry of the Southern North Sea area was recovered from the European Marine Observation and Data Network (EMODNet) data portal^12^ for use in a localized reconstruction. Palaeobathymetry was created by adding isostatic adjustment data^13^ to the bathymetric data obtained from EMODNet using the method identified in Hill et al. (2014)^18^. Small, unresolved islands and features associated with the presence of modern sand banks were removed from all coastlines to aid clarity and consistency. Using the method described by Fruergaard et al.^6^, the local Tsunami height at ELF001A was then extrapolated from the topographic height of the top of the tsunami sequence within the core and applied to the palaeobathymetric data, Figure S1.4. It should be emphasized that the purpose of this method is not to attempt to produce a full or detailed model of the tsunami, but rather to better to generate a visualization to improve our understanding of the spatial nature of this deposit and how it arrived in the basin within which it is situated. To avoid over-fitting, from these data we inferred a more generalized interpretation of the location of the 8.2Kyr coastline and the extent of tsunami run-up across the palaeolandscape by smoothing the line estimates, Figure 2.

The basin itself is an elongated structure trending North-West – South-East. It covers an area of 114 km2, with a flat bottom, gently sloping sides and is 12 meters deep. On the North, NE and NW sides of the basin, the feature is bound by terminal moraines of Late Devensian age, with tunnel valleys forming breaches in these moraine structures^14^. To the south the basin is bound by a slight topographic rise in the Boulders Bank formation. This rise is breached by a Late Devensian outwash structure which was reused in the Early Holocene by a fluvial system, referred to by the project as the ‘Southern River’. Both these breaches in the North and South West form the two main routes for material to run into or out of the basin. Using the information gained from the palaeobathymetry and the localized tsunami height, the way in which the tsunami entered the Southern River system could be ascertained. This was determined to have entered the central basin through the gap in the moraine caused by the glacial tunnel valley and outflowed through the southern section of the channel (Figure 2) was observed. We recovered cores in the southern section that contained the characteristic storm surge deposit, identifying potential tsunami material in ELF0031A, with ELF0039 as a further strong candidate for containing tsunami trace material (figures SF1.1, SF1.3, SF1.4).

**Figure S1.4.**
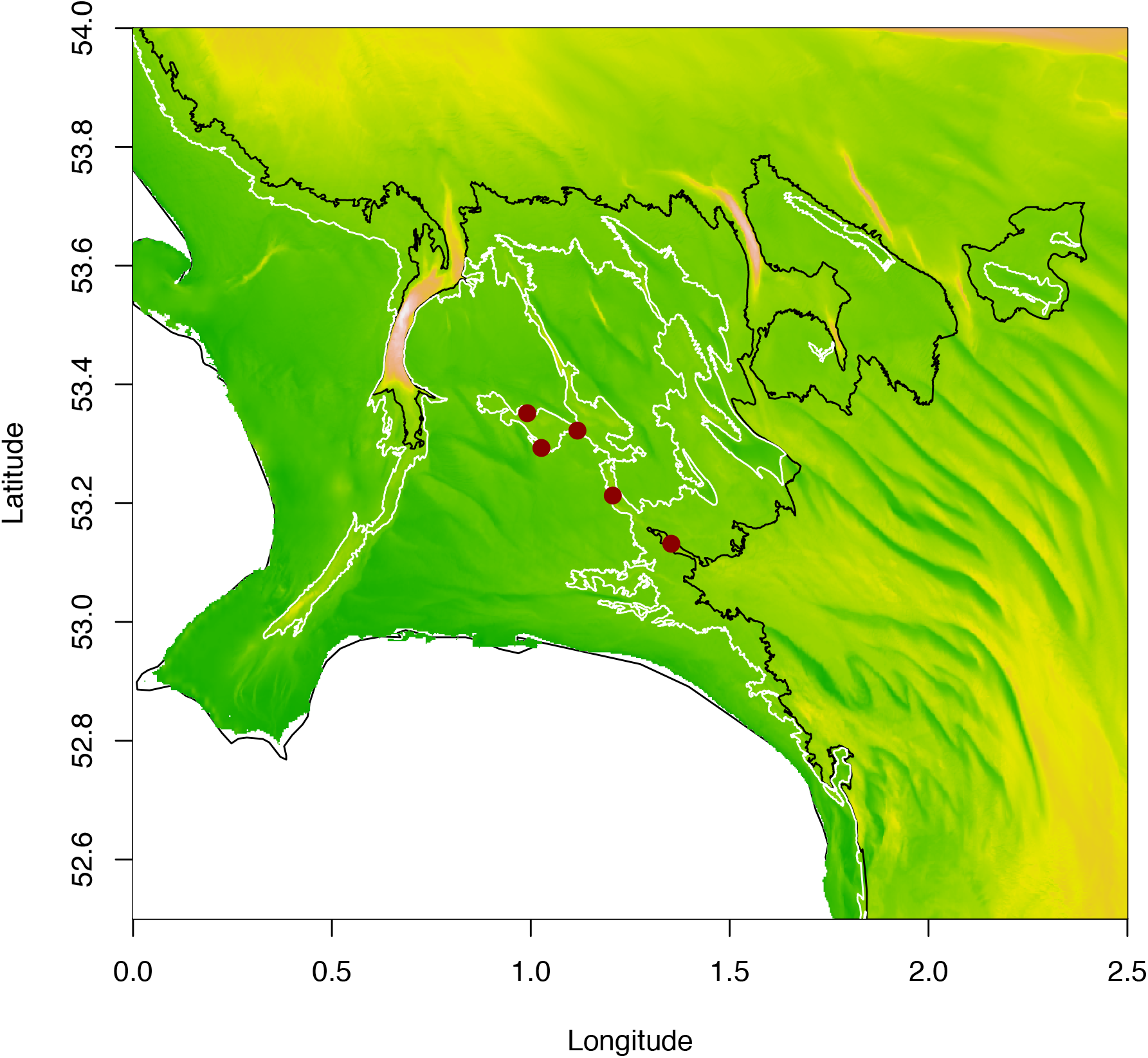
Paleobathymetric estimation of 8.2Kyr coastline and tsunami run-up limit. Coastline indicated in black, tsunami indicated in white. Map coloring indicates relative bathymetric depth. Locations of candidate tsunami associated cores indicated in red circles, from left to right ELF059A, ELF003, ELF001A, ELF031A and ELF039.

### Supplementary Text S2 Geochemistry Analysis

#### S2.1 Palaeomagnetics

sampling of core ELF001A was carried out at the University of Wales Trinity St David Lampeter Campus where the cores are in cold storage. Samples were taken in approximately 0.10m intervals from unit ELF001A-4 at 0.93m; 25 cylindrical samples were obtained (sample diameter 20mm) orientated up core. Samples ELF001a/1, /2 and /3 originate from the disturbed units between 0.90m and 1.51m.

Palaeomagnetic and rock magnetic measurements were carried out at the University of Bradford’s Archaeomagnetic Dating Laboratory and at Lancaster University’s Centre for Environmental Magnetism and Palaeomagnetism (CEMP). At CEMP the 2G Enterprises DC 755 superconducting rock magnetometer with RAPID automatic sample handler, installed in a magnetically shielded room with in-line orthogonal alternating field (AF) demagnetisation was utilised to assess the stability of the natural remanent magnetisation (NRM). All samples were subjected to stepwise AF demagnetisation peak field of 2, 4, 6, 8, 10, 15, 20, 30, 40, 50, 60, 70, 80mT. The 2G RAPID system can generate gyroremanent magnetisation (GRM) by static AF demagnetisation of ferromagnetic materials in fields above ∼30mT. In order to remove this effect, basic measurement procedures set out by Stephenson (1993)^21^ were followed and all results were corrected using the GRM correction tool in the software GM4Edit^22^. After each sample had undergone demagnetisation an anhysteretic remanent magnetisation (ARM) was imparted in a bias DC field of 60µT, 80µT, 100µT, 120µT applied with an in-line single axis DC coil, combined with an alternating field of 80mT. The anhysteretic magnetic susceptibility values were estimated from the slope of linear regression. Subsequently, each sample received a Saturation Isothermal Remanent Magnetisation (SIRM) at 1000mT before applying backfield IRMs at 20mT, 50mT, 100mT, 300mT, and 1000mT. IRMs were applied using a Molspin Ltd Pulse Magnetiser for 20 – 100mT and a Newport electromagnetic for 300 – 1000mT fields.

Magnetic susceptibility in the laboratory was measured using a Bartington MS2 susceptibility meter in addition to the direct measurement of the magnetic susceptibility of the core at 0.05m intervals using a Bartington MS3 susceptibility meter with MS2K attachment. Drift corrections were applied to all measurements of magnetic susceptibility. Reference should be made to Dekkers (2007)^23^ for a thorough discussion on magnetic proxy parameters and their application in palaeomagnetic lacustrine settings^24,25^ where similar protocols were followed. See Walden (1999)^26^ for detailed discussions on ARM and it’s uses as a proxy for relative abundances and magnetic grain sizes.

The magnetic properties of ELF001A are dominated by magnetite with smaller amounts of antiferromagnetic minerals more prevalent at specific horizons within the sediment. The magnetic proxy information calculated from the full suite of analyses shows four distinct magnetic phases indicative of the changing depositional palaeoenvironment.

*0.90 – 1.10m* Corresponding to stratigraphic unit 4 this interval lies above the tsunami event in question. From all the proxy parameters calculated this interval stands out. There appears to be a much larger abundance of magnetic minerals at this horizon as shown by the SIRM and magnetic susceptibility values. The lower percentage of IRM between 0 – 20mT signifies a lower concentration of coarse magnetite than the sediments below. This is also reflected by the ARMχ values showing a higher abundance of ultra-fine magnetite at the Superparamagnetic/Single Domain (SP/SD) boundary^25^. The S-ratio and coercivity of remanence values show a higher concentration of harder magnetic minerals such as haematite and goethite. This represents a significant change in detrital input from the surrounding catchment compared to before the tsunami. The low %FD values (not shown) do not point to a high iron-bearing clay input as would be seen from clay rich soils. Instead the input could be from glacial till which at this time would have been deposited in the coastal area after the glacial retreat. The deposition of glacial till on the surrounding landscape could have weathered to produce haematite, as seen in similar studies of glacial retreat^27^. The increase in the transport of glacial till into this system could result in increasing trends for the magnetic concentration proxies (ARMχ, χlf, and SIRM).

*1.10m – 1.50m* This section relates to stratigraphic unit 6 and is the deposit associated with the tsunami event. Characterised by a much lower χlf value than the surrounding units in parallel to a drop in fine-grained magnetite (ARMχ). This suggests the material brought in by the tsunami is from a different origin to the local sediment supply as it contains a lower abundance of SP particles. The S-ratio for this phase suggests a higher concentration of haematite and goethite, however this is not mirrored by the Hard IRM proxy (not shown) or the coercivity of remanence, suggesting that the magnetic behaviour is the result of multi-domain (MD; coarser grained) magnetite co-existing with fine grained greigite. The presence of greigite particles in this deposit are indicative of intense changes in the palaeoenvironment^25^.

*1.50m – 1.90m* This magnetic phase is distinguished by the %bIRM_0-20mT_ and the coercivity of remanence which show a larger abundance of coarse-grained magnetite up to and including the tsunami interval. At the base of this interval (∼1.90m) the change in these proxies could represent the start of the glacial retreat from the southern North Sea and the gradual sea level rise. The gradual increase in ARMχ reflects the gradual increase in ultra-fine magnetite which would occur as the water column rises in conjunction with fresh glacial till deposition^25,27^.

*1.90m – 3.60m* This phase represents little or no change in detrital input consistent with a low energy estuarine system. The S-ratio gradually increases down this part of the core suggesting a higher ratio of magnetite being deposited through this phase, which could be indicative of a change in salinity.

#### S2.2 Elemental core scan

The surface of each core section of ELF001A was scraped and cleaned to ensure a smooth, flat surface, and the top 3 metres was scanned using an Itrax® XRF core scanner at 500µm resolution with a dwell time of 15 seconds and x-ray tube settings at 30 kV and 50 mA. The modern material in the uppermost core section (Unit ELF001A-1) was not scanned. In individual core sections, the scanning line was adjusted to avoid sampling holes and some sub-sections were run individually to enable the core surface to be kept as flat as possible. Scanning data were compiled to produce a composite sequence. Gaps in the data represent parts of the core which were not scanned. This may have been due to sampling gaps, those instances where the nature of the sediment did not provide a smooth surface for scanning, or the result of poor data quality (low kcps values). Selected elements are presented, normalised to the sum of the incoherent and coherent scattering which account for the effects of Compton scattering and Rayleigh scattering respectively^28–30^.

In general terms, analysis of the ELF001A core suggests that the unit interpreted as indicating a major storm event or tsunami is associated with a rapid switch to marine material. This is indicated by the sulphur signal above the red line in Figure 3. Calcium and strontium signals could derive from terrestrial conditions but these signals differ significantly to the pattern for titanium and potassium which must related to terrestrially derived material. The shaded bar in Figure 3 indicates where the data suggests an increased amount of terrestrial material above the proposed tsunami layer. Bromine may be a proxy for organic matter or sea spray, depending on context. Here it is more likely to represent organic content. Iron follows titanium and does not seem to represent changing redox conditions.

Peaks in the calcium and strontium signals above the layers associated with an enhanced terrestrial signal are further indications for marine conditions and perhaps shells.

#### S2.3 Organic Chemistry profiling

An Agilent 7890A gas chromatograph (GC) coupled with a 5975C Inert XL mass selective detector was used for the lipid analysis. The splitless injector and interface were maintained at 300°C and 340°C respectively. Helium was the carrier gas at constant flow. The temperature of the oven was programed from 50°C (2 min) to 350°C (10 min) at 10°C/min. The GC was fitted with a 30m x 0.25mm, 0.25µm film thickness 5% Phenyl Methyl Siloxane phase fused silica column. The column was directly inserted into the ion source where electron impact (EI) spectra were obtained at 70 eV. Samples were analyzed using a full scan method from *m/z* 50 to 800. For the lipid extraction, fourteen sub-samples from core ELF001A were dried at room temperature for 48 hours, ~3g of each was then solvent extracted using three portions of 12ml (dichloromethane: methanol 2:1 v/v) with ultrasonication and centrifugation. The solvent was transferred into a clean glass vial and removed under a stream of nitrogen at 40°C. The extracts were then silylated with drops of BSTFA at 70°C for an hour. Excess BSTFA was removed under a stream of nitrogen and the samples diluted in 1ml of dichloromethane for analysis.

The lipids analysis of core ELF001A, yielded *n*-alkanes, fatty acids, *n*-alkanol and sterols, of these lipids the *n*-alkanes are the most informative in respect of the origin of the lipids. These show that; the area ELF001A -3 is dominated by marine organic inputs probably from submerged aquatic plants. In area ELF001A -4, aquatic plants are present, with the signals for bacteria and terrestrial plants in significant quantities. In addition, signals of sulfate reducing bacteria were also identified. This area has the chemical profile of an estuarine area, or it may be an area of water present, just before submergence. Area ELF001A -5 has a chemical organic profile similar to that obtained from area ELF001A -3, where submerged marine plants are dominant. In contrast, area ELF001A -6 is the most complex portion with evidence for terrestrial and marine plants and algae within just 15cm of the column, although marine inputs dominate this area. This suggests a major event associated with the deposition of these mixed deposits within well-defined strata. Area ELF001A -7 is the most homogenous of samples examined and is associated with terrestrial plants, bacteria and freshwater within the lower part of the core. The Carbon Preference Index CPI ratio for these samples distinguishes between terrestrial and marine sources. This indicates that terrestrial materials are increasingly present in the lower parts of the core^31,32^ (Figure 3). All NAR ratios are closer to one than zero, which indicates the origin of the lipids from sources other than petroleum^33,34^ (Figure 3).

### Supplementary Text S3 Sediment dating

#### 3.1 OSL dating of sediments

In this supplementary text, we describe the protocols and procedures that were used to determine the quartz SAR OSL ages shown in Table S3.1:

**Table S3.1.**
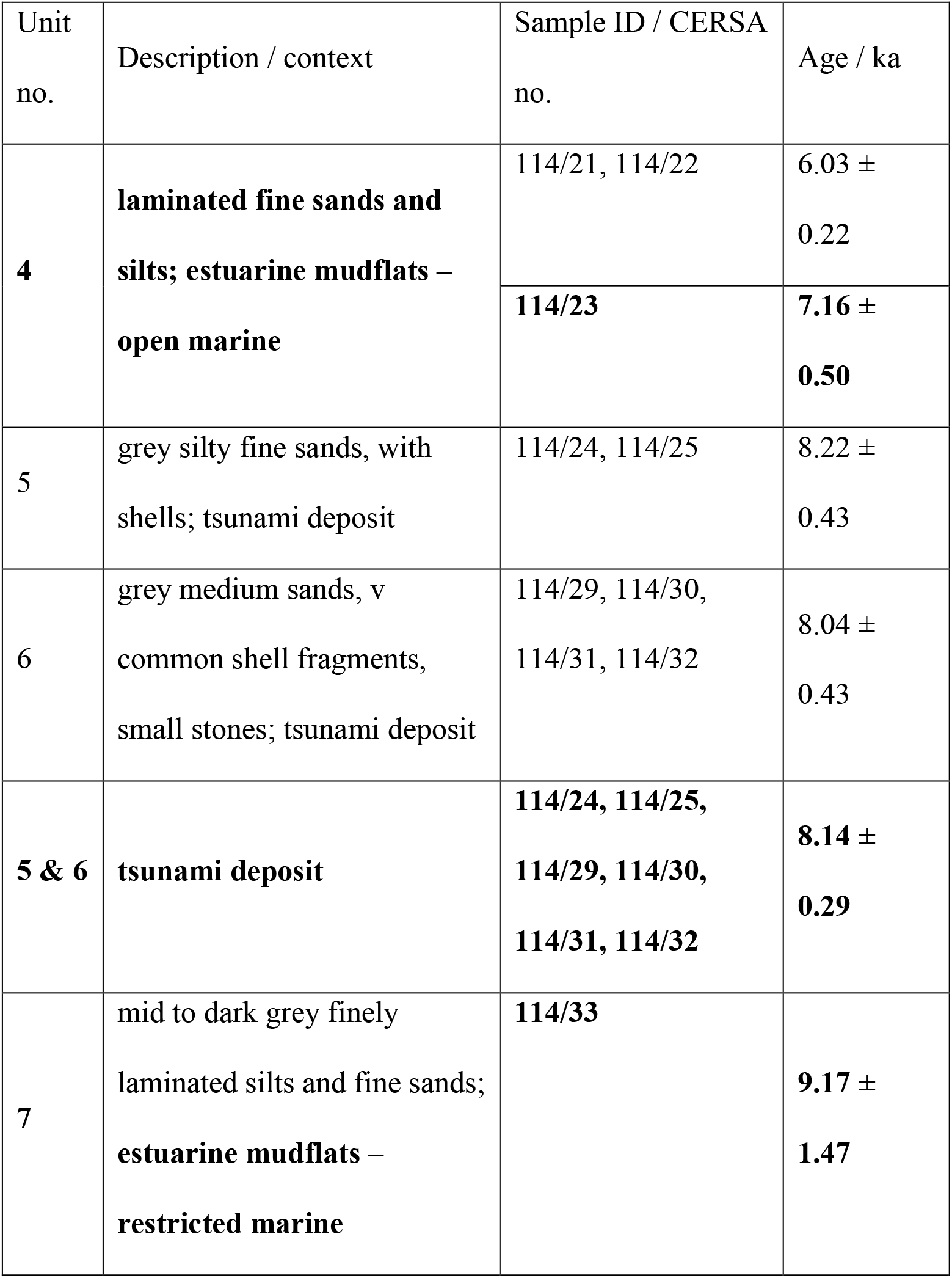
OSL date summary of sediment core ELF001A.

##### A Luminescence screening measurements (Figure S3.1)

The sediments revealed in core ELF001A were first appraised using portable OSL equipment (following procedures reported in Sanderson and Murphy, 2010 and Kinnaird et al., 2017). This allowed for the calculation of IRSL and OSL net signal intensities, their depletion indices and the IRSL - OSL ratio, which were plotted in relation of the lithostratigraphy of the core (Figure S1.1). This proxy information was used to select the most promising intervals in the core for dating: at the base of the *open marine* estuarine mudflats (unit 4), through the tsunami deposit (units 5 and 6), and at the top of the *restricted marine* estuarine mudflats (unit 7).

##### B. Calibrated luminescence measurements (figures S3.1 and S3.2)

Calibrated luminescence screening methods, as previously utilised by Kinnaird et al. (2017)^35^ were used to generate stored sensitivity- and dose-depth profiles for core ELF001A. Luminescence sensitivities (photon counts per Gy) and stored dose (Gy) were evaluated on paired aliquots of HF-etched quartz, using procedures modified from Burbidge et al. (2007)^37^, Sanderson et al. (2003)^38^ and Kinnaird et al. (2017)^35^. This calibrated dataset is shown relative to the proxy information and the lithostratigraphy in Figure S3.1.

All OSL measurements were carried out using either Risø TL/OSL DA-20 or DA-15 automated dating systems, equipped with a ^90^Sr/^90^Y β-source for irradiation (dose rates at time of measurement, 1.10 and 0.03 Gy/s, respectively), blue LEDs emitting around 470 nm and infrared diodes emitting around 830 nm for optical stimulation. OSL was detected through 7.5 mm of Huoya U-340 filter and using a 9635QA photomultiplier tube. OSL was measured at 125°C for 60 s. The OSL signals, *Ln* and *Lx*, used for equivalent dose (*De*) determinations were obtained by integrating the OSL counts in the first 2.4 s and subtracting an equivalent signal taken from the last 9.6 s. The protocol implemented here involved a readout of the natural signal, followed by a 1 Gy test dose, then readouts of the regenerated cycles following a series of nominal doses between 5 and 120 Gy, each with a subsequent 1 Gy test dose. For all, OSL followed a preheat of 220°C and was measured at 125°C for 60s.

Apparent dose estimates were made in Luminescence Analyst v.4.31.9, using dose response curves forced through zero and the two to three normalized regenerative points with an exponential function (Figure S3.2).

##### C. Equivalent dose determinations (figures S3.3 to S3.5)

Samples selected for full quantitative quartz OSL dating were subjected to further mineral purification of quartz (cf. Kinnaird et al. 2017)^35^. Equivalent doses were determined by OSL on at least 20 aliquots per sample (typically 40+ aliquots) using a single aliquot regenerative dose (SAR) OSL protocol (cf. Murray and Wintle, 2000^39^; Kinnaird et al. 2017^35^; see S3 therein). This was implemented, using five regenerative doses (nominal doses between 1 and 40 Gy), with additional cycles for zero dose, repeat or ‘recycling’ dose (2.5 Gy) and IRSL dose (2.5 Gy). Five preheat temperatures were explored between 220 and 260°C, in 10°C increments.

Data reduction and De determinations were made in Luminescence Analyst v.4.31.9. Individual decay curves were scrutinised for shape and consistency. Dose response curves were fitted with an exponential function, with the growth curve fitted through zero and the repeat recycling points. Error analysis was determined by Monte Carlo Stimulation.

The equivalent dose distributions for each sample are shown relative to the lithostratigraphy of the core in figures S3.3 and S3.4, for the grain size fractions 90-150µm and 150-250µm (see also tables S3.2 and S3.3). Individual sample distributions were appraised for equivalent dose homogeneity, and, when the luminescence profiles suggested stratigraphic coherence, different combinations of merged datasets across stratigraphic associations were explored (e.g. Fig. S3.5). Different permutations of the assimilation of equivalent doses to obtain the burial dose were considered, including weighted combinations and statistical dose models^40^. It was concluded that, with the dosimetry as presently constrained, the combined distributions were most appropriate for calculation of the stored dose. In justification of this, the stored doses thus obtained correlate well with the apparent dose-depth profiles obtained earlier (R^2^ = 0.943).

##### D. Dose rate determinations (Figure S3.6)

The dose rates to these materials were assessed through a combination of X-ray Fluorescence core scanning, high-resolution gamma spectrometry (HRGS), and inductively-coupled plasma mass spectrometry (ICPMS) analysis.

Semi-quantitative element concentrations of K, U and TH, as obtained by X-ray Fluorescence core scanning at Aberystwyth University are shown in Figure S3.6.

HRGS measurements were performed at the Environmental Radioactivity Laboratory (ERL; UKAS Testing Lab 2751), University of Stirling. All sample handling, processing and analysis were undertaken in accordance, and in compliance with ERL protocols LS03.1, 03.2 & 03.6 and LS08. HRGS measurements were performed on a High Purity Germanium detector. Standard laboratory efficiency calibrations were used, derived from GE Healthcare Ltd QCY48 Mixed Radionuclide Spike and DKD RBZ-B44 ^210^Pb spike. All absolute efficiency calibrations were corrected for variations in sample density and matrix. HRGS measurements were undertaken on composite bulk sediment samples at 108cm, 150cm, 155cm and 360cm.

Radionuclide concentrations determined by HRGS are listed in Table S3.4.

Concentrations of K, U, Th and Rb were measured directly using ICPMS at the STAiG istope labs at the University of St Andrews. ICPMS measurements were performed on dried, homogenised sub-samples of sediment taken from discrete horizons in the core. 15-20 g of sediment were taken from each sample, then ground and homogenised using a Tema Machinery Disc Mill. 2 gram sub-samples were treated in a furnace set at 1000 °C for 6 hours. 50 mg quantities of sediment from each sub-sample were prepared for ICPMS using total rock digestion by Ammonium Bifluoride^41^, adapted to include additional fluxes in hot HCl and HNO_3_. Samples were prepared by gravimetric serial dilution at 10 and 1000x in 0.4 M HNO3:0.02 M HF, for analysis of U and Th, and K, respectively. All analyses were conducted on an Aligent 7500 ICP-MS instrument. Samples and standards were introduced through a PFA spray chamber in 0.4 M HNO3:0.02 M HF using a self-aspirating nebuliser (100 µL min-1). Analytical calibration standards (0.1, 1, 10, 100 and 500 ppb for all elements) were prepared by gravimetric serial dilution from 10 ppm certified stock multi-element solutions (Agilent) in 0.4 M HNO3:0.02 M HF. Inter-calibration between counting and analogue detection modes was performed prior to each analytical session. ICPMS analyses were performed on samples CERSA119 [95-100cm depth], 114/21-22 [100cm], 120 [100-105cm], 122 [110-117cm], 114/28-29 [129-136cm], 114/31-32 [146-150cm], 114/33-34 [155-160cm], 114/35-36 [165-170cm] and ELF114/4-45 [201-211cm]. An IAG reference material, SdAR-L2 blended sediment, was prepared and run in batch with the ELF001A core samples. Analytical calibration standards (0.1, 1, 10 and 100 ppb for all elements) were prepared by gravimetric serial dilution from 10 ppm certified stock multi-element solutions (Agilent) in 0.4 M HNO3:0.02 M HF. Additional calibration standards at 2 and 75 ppb were prepared and run as ‘samples’ within the sample batch.

Radionuclide concentrations determined by ICPMS are listed in Table S3.5.

These data were used to determine infinite matrix dose rates for α, γ and β radiation, using the conversion factors of Guérin et al. (2011)^42^, grain-size attenuation factors of Mejdahl (1979)^43^ and attenuated for sediment-matrix water contents. Table S3.6 lists the effective beta, gamma and total environmental dose rates to HF-etched, 90-150µm quartz.

#### E. Age determinations

Luminescence ages are calculated as the quotient of the stored dose (or burial dose, Gy section C above) and the environmental dose rate to these materials (mGy a^−1^; section D; Table S3.7). The resolution at which the stored doses were constrained (at 5 to 10 cm through the tsunami deposit) is not matched by the resolution at which the dosimetry is constrained (>15 cm). From the core scan it is known that there are significant variations in K concentrations within the tsunami deposits, with positive and negative gradients and also spikes. It was thus concluded that combining equivalent doses from each of the units - 4, 5, 6 and 7 – provided the most appropriate method of assimilation for calculating depositional ages. The results of these determinations support a link to the Storegga tsunami and are coincident with dating for this event between 8120 -8175BP, and provided through other studies^44–46^.

**Figure S3.1:**
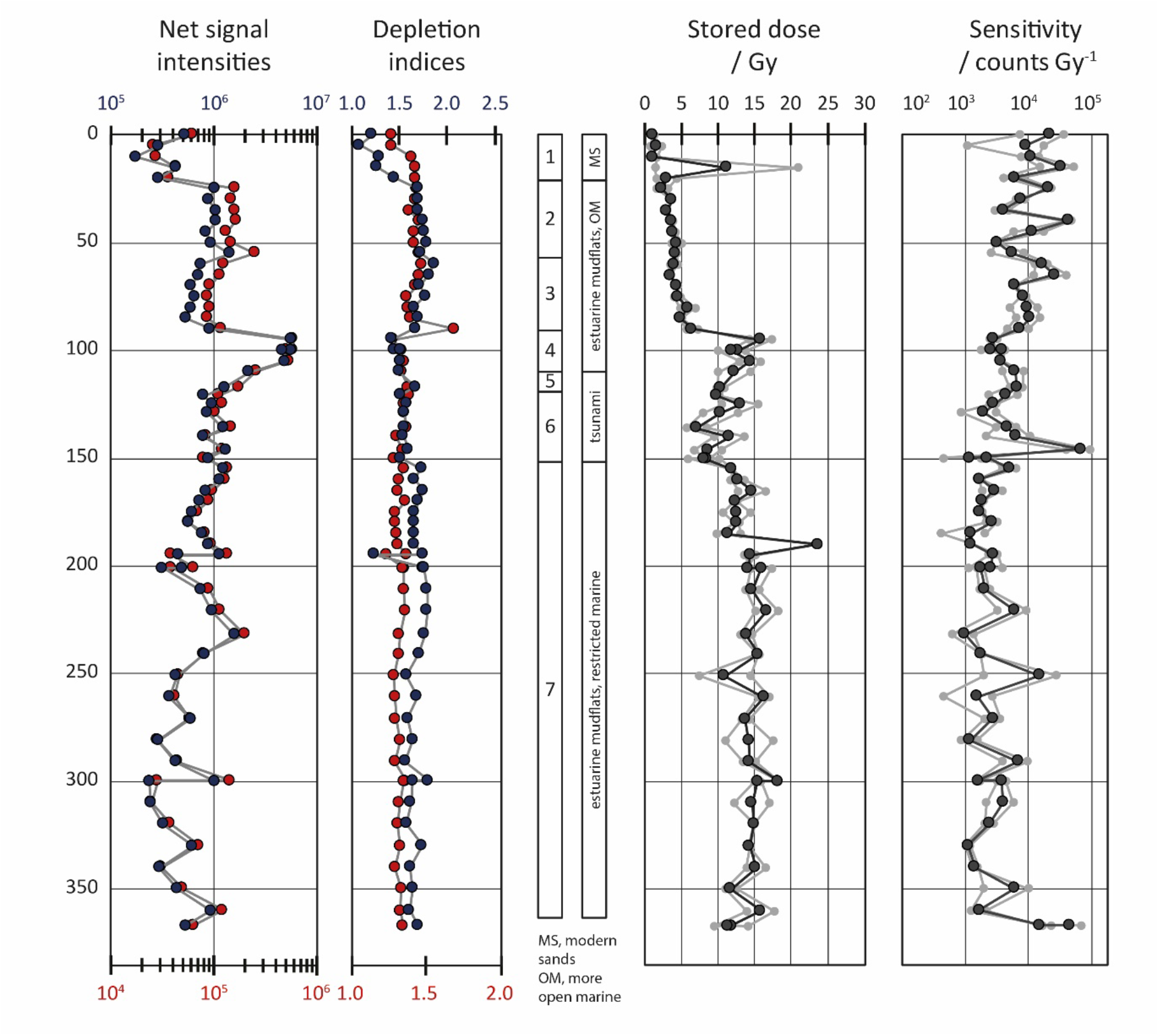
Luminescence stratigraphies for ELF001A: (left) proxy luminescence-depth profile generated at sampling using portable OSL equipment, (right) stored dose- and sensitivity-depth profiles based on a simplified SAR OSL on paired aliquots of HF-etched quartz

**Figure S3.2:**
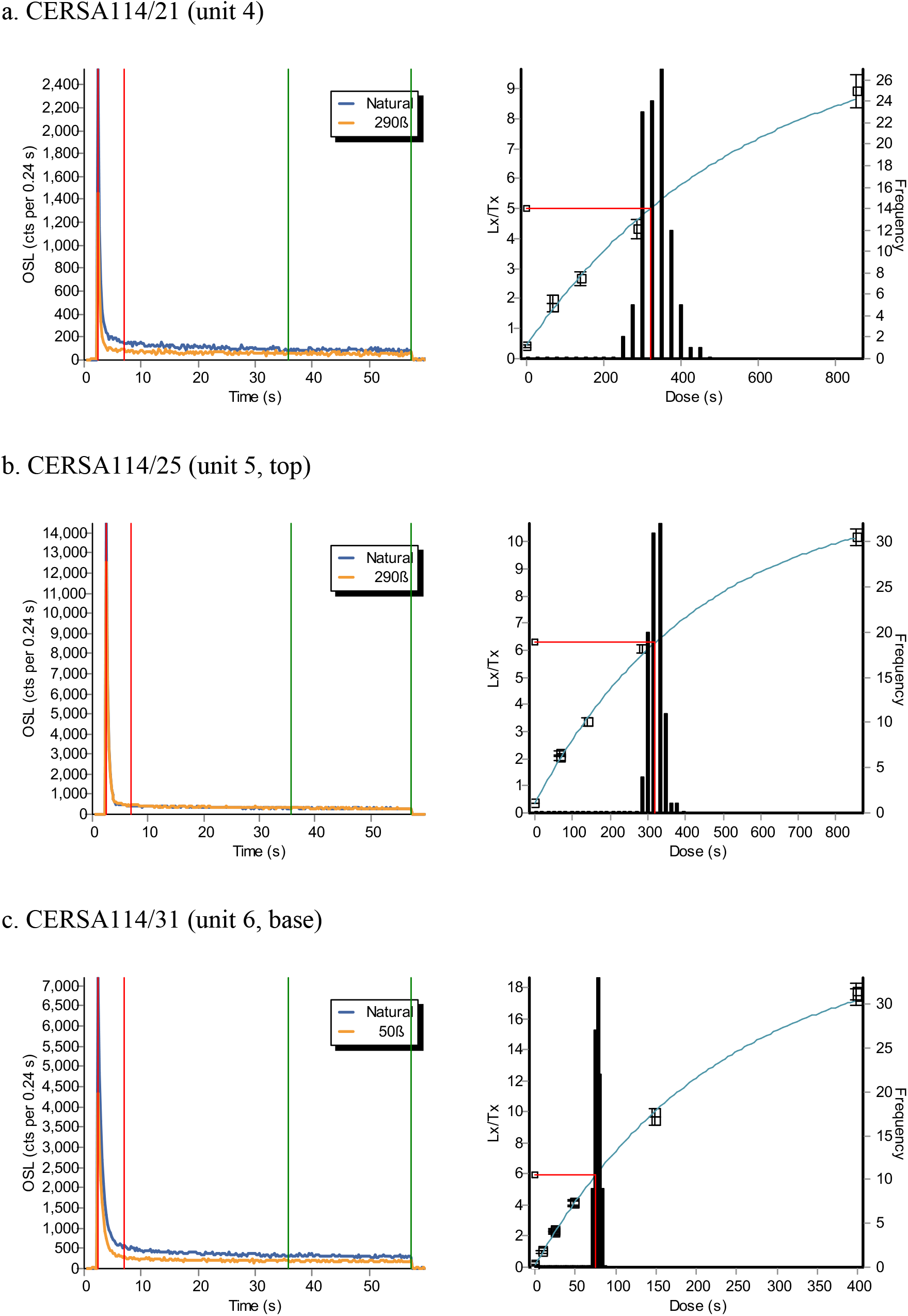
Representative decay (left) and composite dose (right) curves.

**Figure S3.3:**
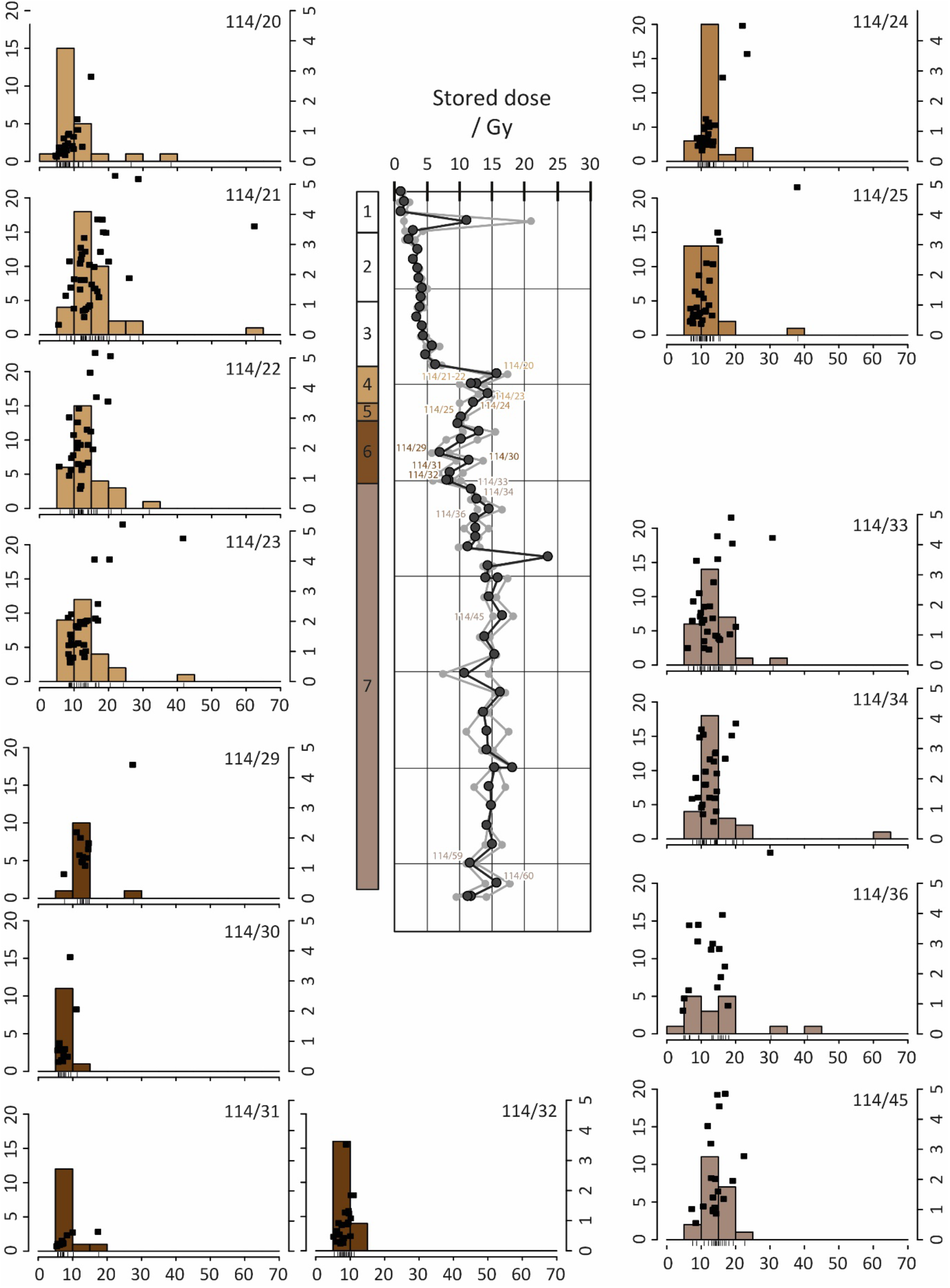
Core ELF1A, De estimates plotted vs depth in the stratigraphy, for 90-150micron quartz.

**Figure S3.4:**
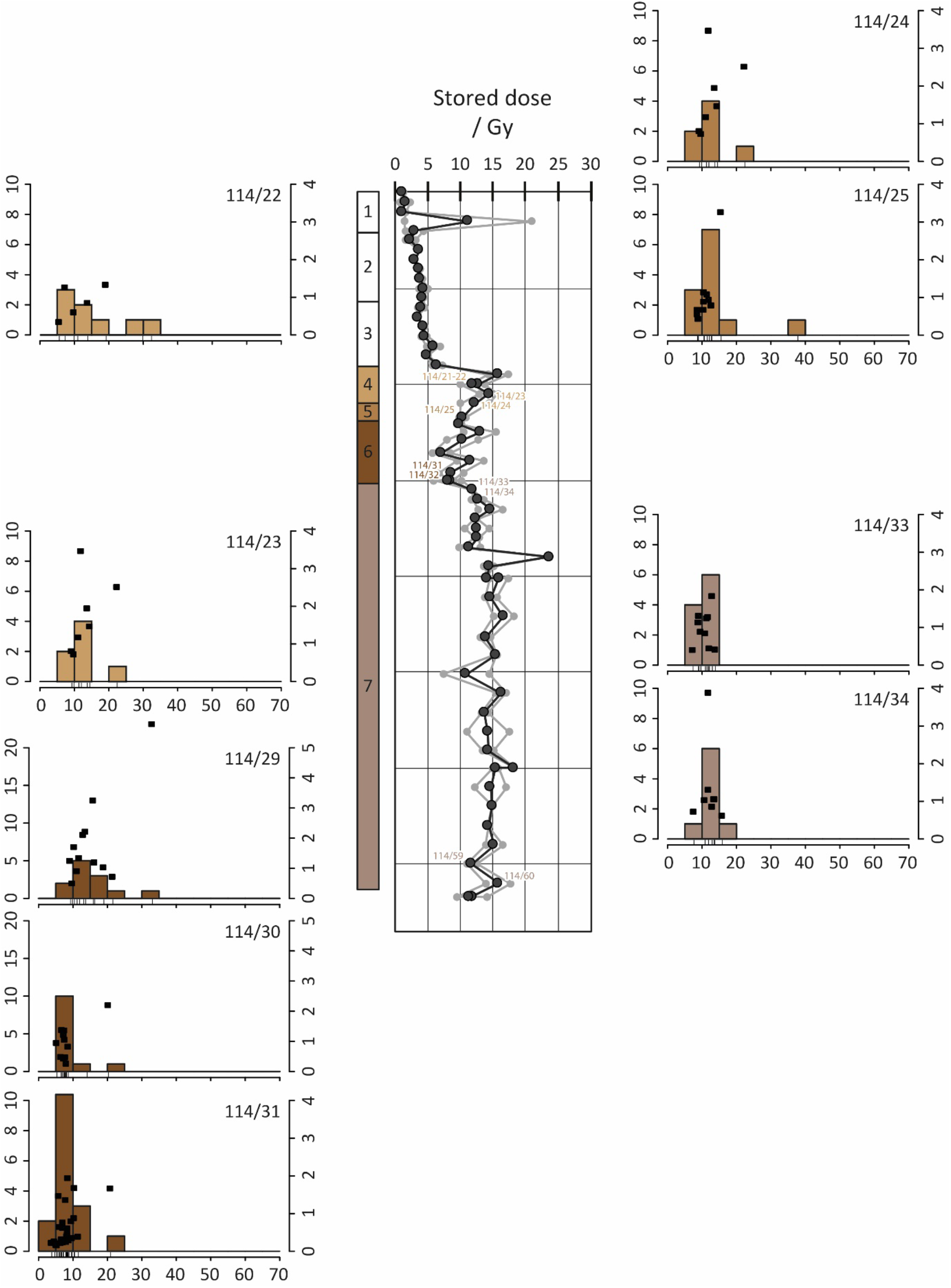
Core ELF1A, De estimates plotted vs depth in the stratigraphy, for 150-250micron quartz.

**Figure S3.5:**
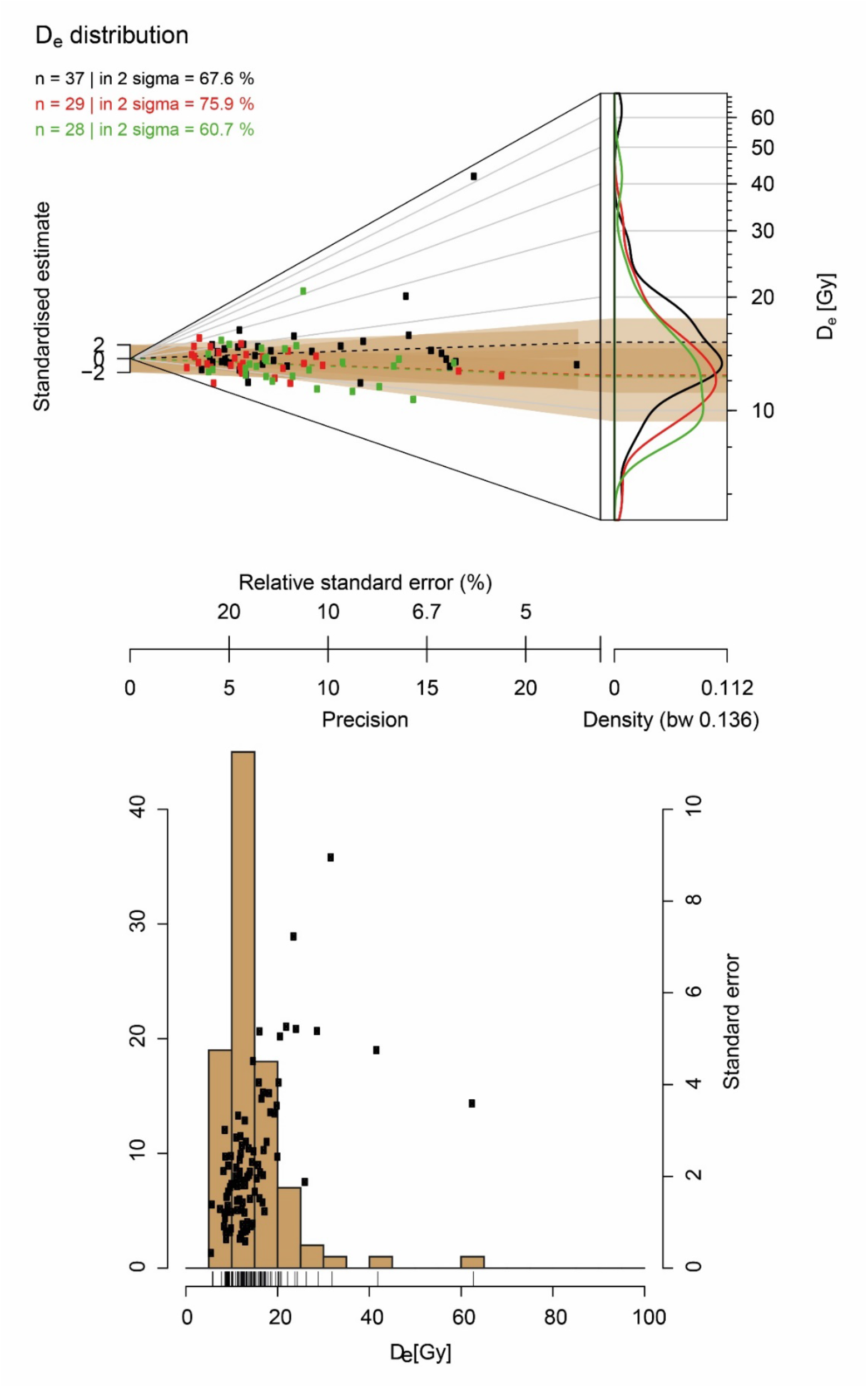
Dose distribution analysis as applied to ELF001A. Unit 4 (tidal mudflats), illustrated as an Abanico Plot and Histogram plot (generated with R luminescence package^47^.

**Figure S3.6:**
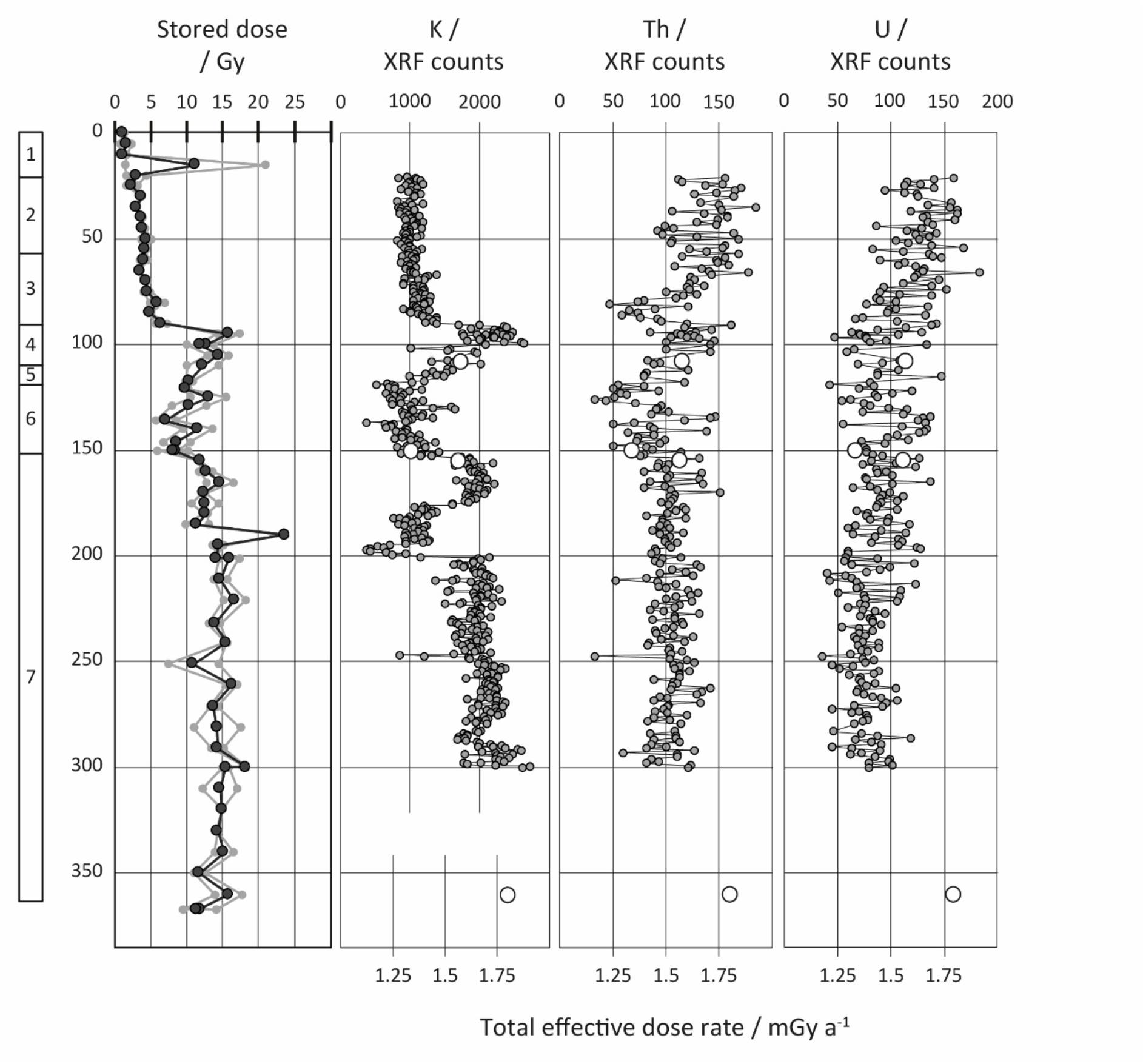
Core ELF001A, semi-quantitative element concentrations for K, U and Th. Obtained by X-ray Fluorescence using the Itrax® core scanner at Aberystwyth University. Also shown, are the total effective dose rates as estimated from HRGS

**Table S3.2:**
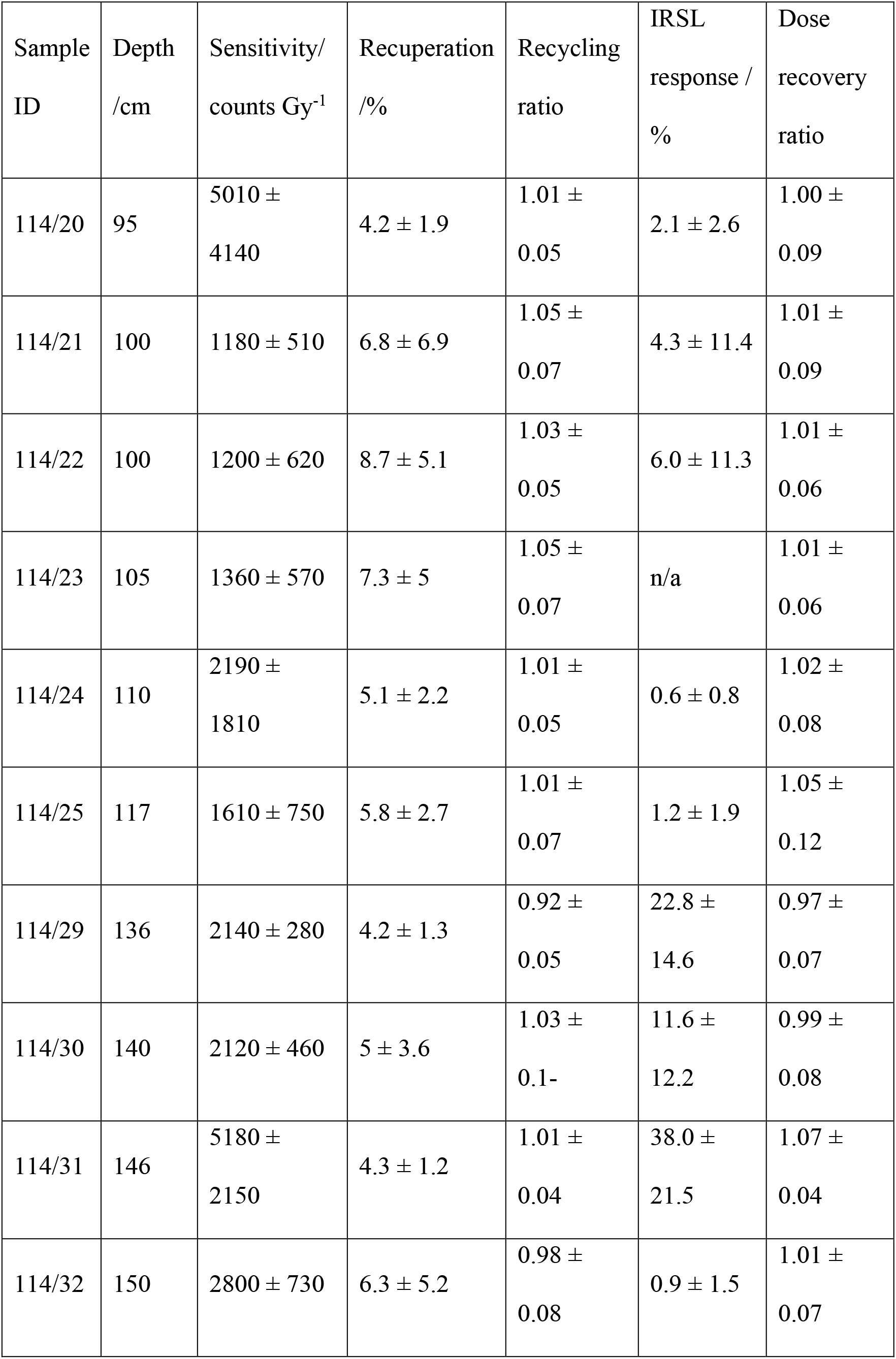
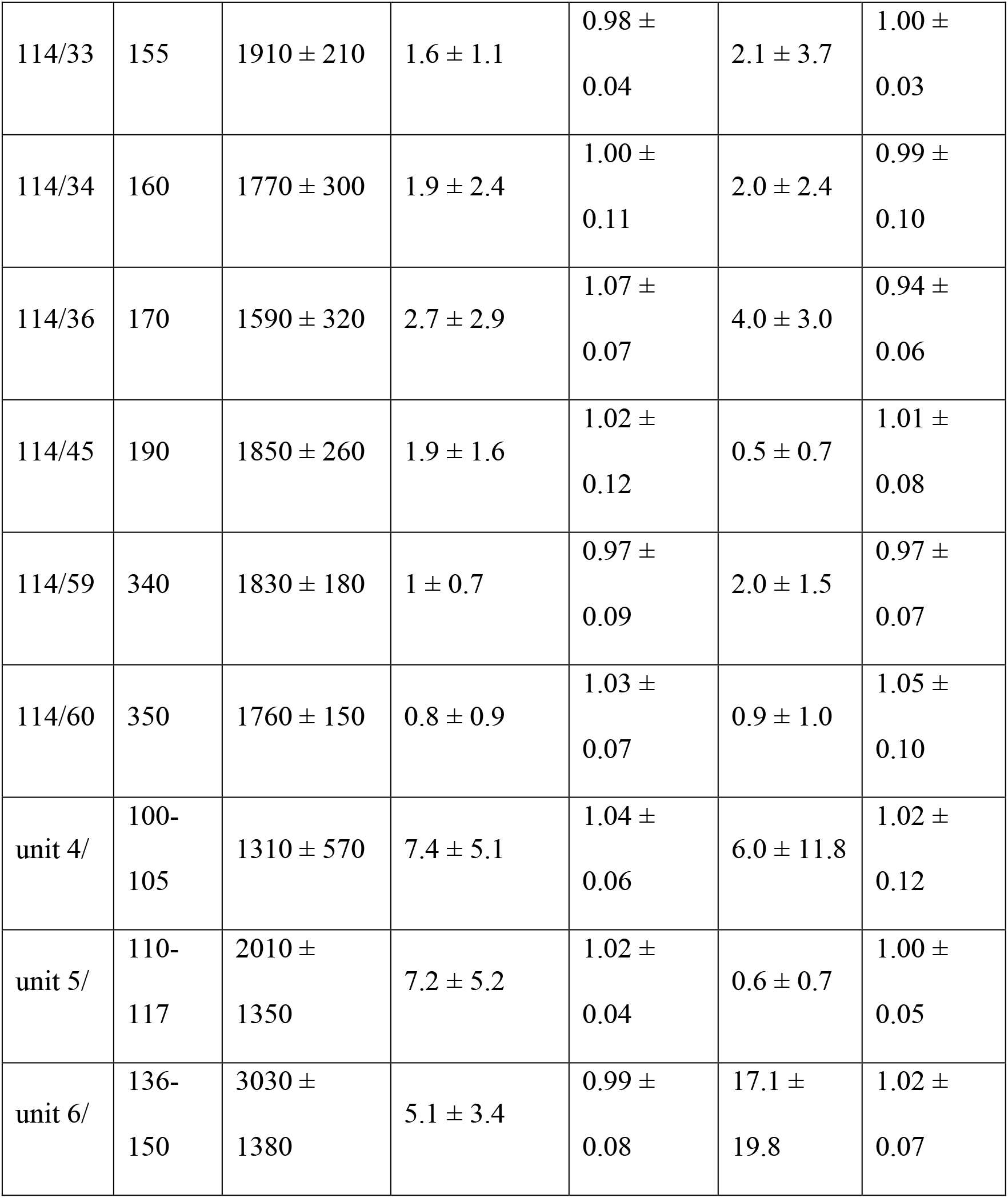
SAR quality criteria, 90-150µm quartz.

**Table S3.3:** SAR quality criteria, 150-250 µm quartz.

| Sample ID | Depth /cm | Sensitivity/<br>counts Gy<sup>-1</sup> | Recuperation /% | Recycling ratio | IRSL response / % | Dose recovery ratio |
| --- | --- | --- | --- | --- | --- | --- |
| 114/20 | 95 | n/a |  |  |  |  |
| 114/21 | 100 | n/a |  |  |  |  |
| 114/22 | 100 | 1390 ± 810 | 6.3 ± 1.5 | 1.05 ± 0.07 | 0.48 ± 0.46 | 1.00 ± 0.08 |
| 114/23 | 105 | 2090 ± 1000 | 6.8 ± 6.2 | 1.06 ± 0.06 | n/a | 0.94 ± 0.01 |
| 114/24 | 110 | 3690 ± 1400 | 5.3 ± 2.8 | 1.00 ± 0.03 | 2.8 ± 3.1 | 1.02 ± 0.05 |
| 114/25 | 117 | 3260 ± 1580 | 3.0 ± 2.7 | 0.99 ± 0.05 | 2.3 ± 3.4 | 1.02 ± 0.07 |
| 114/29 | 136 | 2310 ± 500 | 3.4 ± 2.5 | 0.97 ± 0.11 | 17.0 ± 18.6 | 0.96 ± 0.09 |
| 114/30 | 140 | 2560 ± 560 | 8.4 ± 6.7 | 1.02 ± 0.06 | 3.6 ± 5.6 | 0.99 ± 0.05 |
| 114/31 | 146 | 6620 ± 5820 | 3.8 ± 1.3 | 0.98 ± 0.04 | 36.9 ± 23.8 | 1.07 ± 0.06 |
| 114/32 | 150 | n/a |  |  |  |  |
| 114/33 | 155 | 2550 ± 420 | 3.8 ± 3.6 | 1.06 ± 0.05 | 0.62 ± 0.70 | 0.99 ± 0.06 |
| 114/34 | 160 | 1850 ± 360 | 4.1 ± 3.2 | 0.98 ± 0.07 | 1.3 ± 1.5 | 0.95 ± 0.10 |
| 114/36 | 170 | n/a |  |  |  |  |
| 114/45 | 190 | n/a |  |  |  |  |
| 114/59 | 340 | n/a |  |  |  |  |
| 114/60 | 350 | n/a |  |  |  |  |
| unit 4 | 100-105 | $1740 \pm 900$ | $6.6 \pm 4$ | $1.05 \pm$<br>0.05 | $0.48 \pm$<br>0.46 | $0.97 \pm$<br>0.06 |
| unit 5 | 110-117 | $3350 \pm$<br>1230 | $8.4 \pm 5$ | $1.01 \pm$<br>0.05 | $1.6 \pm 2.4$ | $1.00 \pm$<br>0.05 |
| unit 6 | 136-150 | $4100 \pm$<br>2920 | $4.7 \pm 4.3$ | $0.97 \pm$<br>0.10 | $24.6 \pm$<br>24.2 | $1.01 \pm$<br>0.06 |

**Table S3.4:**
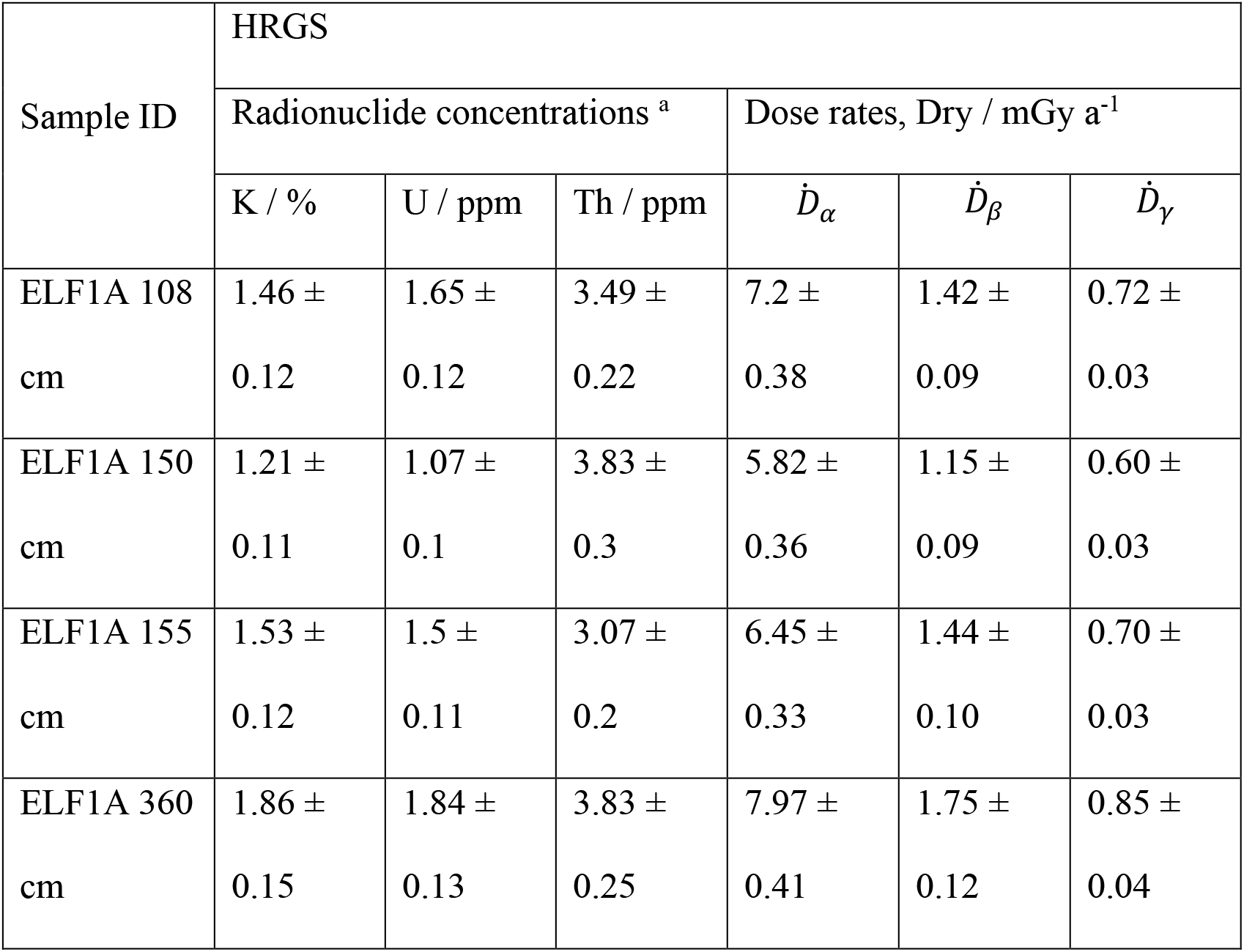
Radionuclide concentrations as determined by HRGS, converted to dry dose rates using the conversion factors of Guérin et al. (2011)

**Table S3.5:** Radionuclide concentrations as directly measured by ICPMS.

| Sample ID | Equivalent<br>to | Depth<br>/cm | K / % | U / ppm | Th / ppm |
| --- | --- | --- | --- | --- | --- |
| 119 [95-100] | 114/20 | 95-100 | $1.75 \pm 0.07$ | $2.22 \pm 0.12$ | $8.2 \pm 0.4$ |
| 114/21-22 [100] | 114/21-22 | 100 | $2.52 \pm 0.06$ | $3.63 \pm 0.18$ | $14.9 \pm 0.8$ |
| 120 [100-105] | 114/21-22 | 100-105 | $1.95 \pm 0.05$ | $2.53 \pm 0.09$ | $9.1 \pm 0.3$ |
| 122 [110-117] | 114/24-25 | 110-117 | $1.37 \pm 0.05$ | $1.84 \pm 0.09$ | $5.6 \pm 0.3$ |
| 114/28-29 [129-136] | 114/28-29 | 129-136 | $0.81 \pm 0.03$ | $1.24 \pm 0.01$ | $4.3 \pm 0.2$ |
| 114/31-32 [146-150] | 114/31 | 146-150 | $0.82 \pm 0.02$ | $0.88 \pm 0.04$ | $2.5 \pm 0.1$ |
| 114/33-34 [155-160] | 114/33 | 155-160 | $2.27 \pm 0.04$ | $2.73 \pm 0.08$ | $10.5 \pm 0.3$ |
| 114/35-36 [165-170] | 114/34 | 165-170 | $2.23 \pm 0.06$ | $2.44 \pm 0.06$ | $9.3 \pm 0.3$ |
| 114/44-45 [201-211] | 114/40 | 201-211 | $2.24 \pm 0.10$ | $2.37 \pm 0.03$ | $8.8 \pm 0.1$ |
| SdAR-L2 | - | - | $3.57 \pm 0.18$ | $2.04 \pm 0.10$ | $24.3 \pm 1.2$ |

**Table S3.6:** Environmental dose rates to HF-etched quartz, reconciled from the HRGS and ICPMS data, and attenuated for grain size and water content.

| Sample ID | Depth /cm | Effective beta dose rate / mGy a<sup>-1</sup> | Effective gamma dose rate / mGy a<sup>-1</sup> | Total effective environmental dose rate / mGy a<sup>-1</sup> |
| --- | --- | --- | --- | --- |
| 114/20 | 95 | 1.30 ± 0.07 | 0.69 ± 0.03 | 2.00 ± 0.05 |
| 114/21 | 100 | 1.98 ± 0.10 | 0.68 ± 0.04 | 2.66 ± 0.08 |
| 114/22 | 100 | 1.46 ± 0.07 | 0.63 ± 0.04 | 2.09 ± 0.06 |
| 114/23 | 105 | 1.18 ± 0.11 | 0.57 ± 0.05 | 1.75 ± 0.07 |
| 114/24 | 110 | 1.02 ± 0.06 | 0.44 ± 0.07 | 1.46 ± 0.08 |
| 114/25 | 117 | 0.63 ± 0.03 | 0.42 ± 0.07 | 1.05 ± 0.07 |
| 114/29 | 136 | 0.59 ± 0.06 | 0.42 ± 0.07 | 1.01 ± 0.07 |
| 114/30 | 140 | 0.59 ± 0.06 | 0.42 ± 0.07 | 1.00 ± 0.07 |
| 114/31 | 146 | 0.57 ± 0.03 | 0.42 ± 0.07 | 0.99 ± 0.07 |
| 114/32 | 150 | 1.68 ± 0.08 | 0.62 ± 0.06 | 1.00 ± 0.13 |
| 114/33 | 155 | 1.64 ± 0.15 | 0.66 ± 0.05 | 1.65 ± 0.18 |

**Table S3.7:**
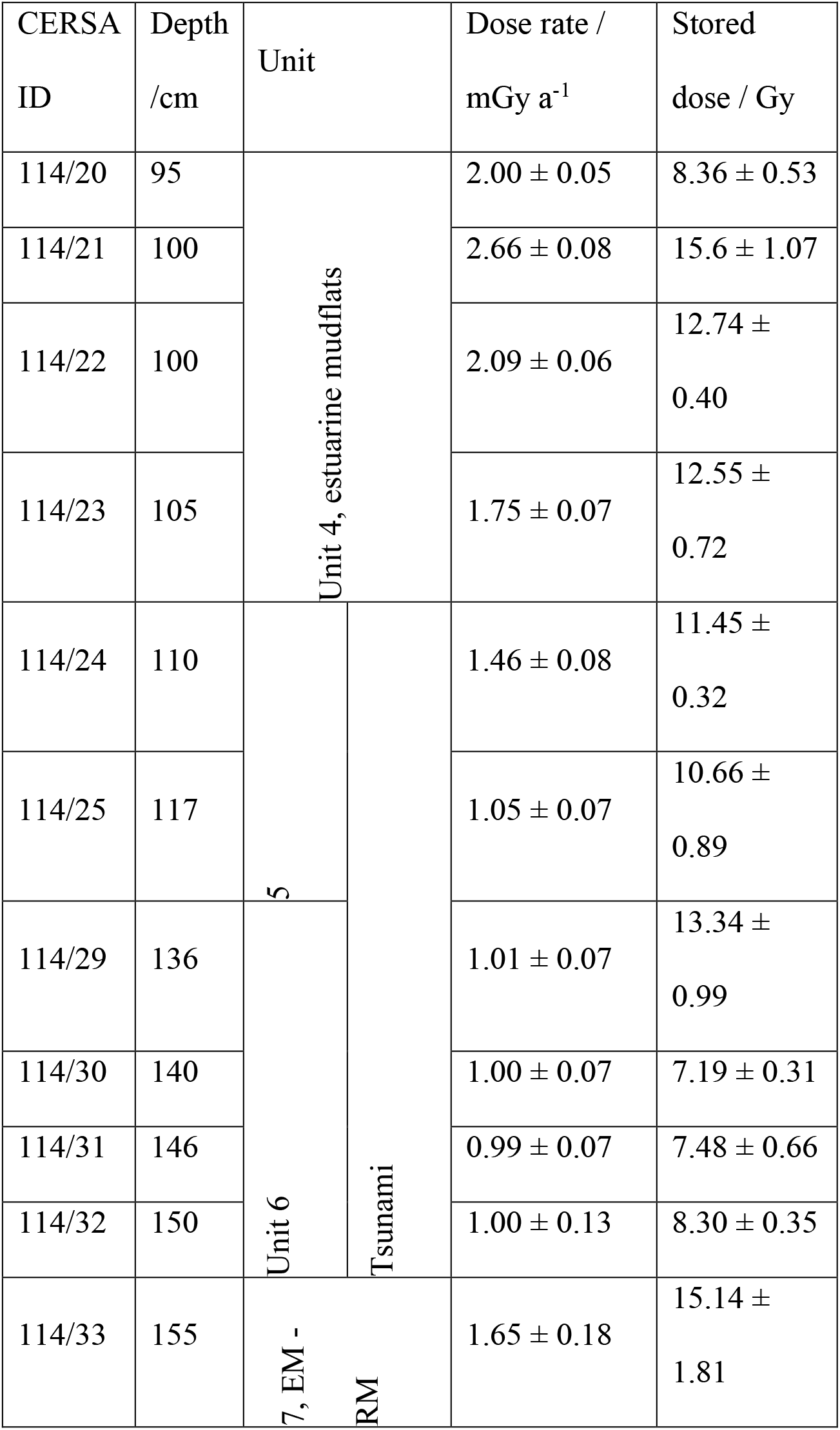
Dose rates and stored doses for the 90-150 µm fraction for ELF001A.

| CERSA ID | Depth /cm | Unit |  | Dose rate / mGy a<sup>-1</sup> | Stored dose / Gy |
| --- | --- | --- | --- | --- | --- |
| 114/20 | 95 | Unit 4, estuarine mudflats |  | 2.00 ± 0.05 | 8.36 ± 0.53 |
| 114/21 | 100 |  |  | 2.66 ± 0.08 | 15.6 ± 1.07 |
| 114/22 | 100 |  |  | 2.09 ± 0.06 | 12.74 ± 0.40 |
| 114/23 | 105 |  |  | 1.75 ± 0.07 | 12.55 ± 0.72 |
| 114/24 | 110 | 5 | Tsunami | 1.46 ± 0.08 | 11.45 ± 0.32 |
| 114/25 | 117 |  |  | 1.05 ± 0.07 | 10.66 ± 0.89 |
| 114/29 | 136 | Unit 6 |  | 1.01 ± 0.07 | 13.34 ± 0.99 |
| 114/30 | 140 |  |  | 1.00 ± 0.07 | 7.19 ± 0.31 |
| 114/31 | 146 |  |  | 0.99 ± 0.07 | 7.48 ± 0.66 |
| 114/32 | 150 |  |  | 1.00 ± 0.13 | 8.30 ± 0.35 |
| 114/33 | 155 | 7, EM - RM |  | 1.65 ± 0.18 | 15.14 ± 1.81 |

##### S3.2 AMS Radiocarbon dating

Radiocarbon measurements were taken on shells from molluscs recovered from 140–145 cm depth in ELF001A, 211-212cm and 314-316cm depth in ELF003, Table S3.8. The shells were submitted to Beta Analytic Inc. where they were pretreated following methods found on their website (https://www.radiocarbon.com/carbon-dating-shells.htm) and dated by accelerator mass spectrometry (AMS). Beta Analytic round all uncalibrated radiocarbon ages to the nearest 10 years, according to the Trondheim convention^48,49^ and assign a conservative minimum error of ±30 ^14^C years.

The radiocarbon age is calibrated in OxCal v4.3^50^ using the internationally agreed Marine13 calibration curve of Reimer et al (2013)^51^ and a local marine reservoir correction (ΔR) of −3 ±99 years, which was calculated using the 14Chrono Centre database (http://calib.org/marine/) and the 10 nearest data points to 55.1369’ N, 3.4086’ W.

**Table S3.8.** Calibrated radiocarbon dates from tsunami associated strata in cores ELF001A and ELF003.

| Core | substrate | calibration | depth (cm) | Age (BP) | range +/- |
| --- | --- | --- | --- | --- | --- |
| ELF001A | Shell | Marine13 | 140-145 | 8340 | 30 |
| ELF003 | Shell | Marine13 | 211-212 | 8400 | 30 |
| ELF003 | Shell | Marine13 | 314-316 | 8350 | 30 |

### Supplementary Text 4 Palaeoenvironmental proxy analyses

#### S4.1 Foraminifera and ostracods

A rapid assessment of the samples was undertaken on 15 samples. A range of materials were present in the samples including plant debris and seeds, molluscs, diatoms, and insect remains. Foraminifera and ostracods were present in all samples.

Three microfossil facies associations have been identified from the samples (Table S4.1). The lowermost facies (associated with unit ELF001A-7) appears to be one indicative of estuarine mudflats. The microfaunas are very restricted (suggesting brackish conditions). The foraminifera are often very small and the ostracods are represented invariably by small juveniles. This is odd and may be, in part, a function of reduced salinities in the environment. This lower part of the sequence also contains the remains of many spirorbid polychaete worms, which are normally attached either to a hard substrate or seaweed. In the absence of a hard substrate it is likely that seaweed was common and this suggests an abundance of algae on the mudflats. There are also a few juvenile molluscs in this part of the sequence. The “marine” component is also limited and therefore suggests the site is open to the estuary or part of a gulf (with reduced salinities). The basal sample also contains rare *Jadammina macrescens* which is a high saltmarsh foraminifer and may be indicative of saltmarsh in the vicinity of the sampling site at this time. Its disappearance above 3.67m may suggest waning saltmarsh conditions.

The second facies type is associated with the coarser sediments of units EF001A-5 and ELF001A-6 that contains both brackish hydrobids and marine oyster shells. These remains are very fragmentary and typically smashed. The ostracods, especially the brackish component, also exhibits damage with many valves broken. This part of the sequence contains a greater number of outer estuarine and marine forams alongside the tidal mudflat and estuarine foraminifera and ostracods noted below. Unit EF001A-5 also contains a few agglutinating foraminifera of high saltmarsh (*Trochimmina inflata* and *Jadammina macrescens*).

The third facies type is associated with unit ELF001A-4 and is dominated by estuarine mudflat species. By comparison with the lower mudflat facies this association appears to be indicative of more open estuarine conditions evidenced by more diverse microfaunas especially of adult brackish ostracods and a greater outer estuarine marine component in general.

**Table S4.1.**
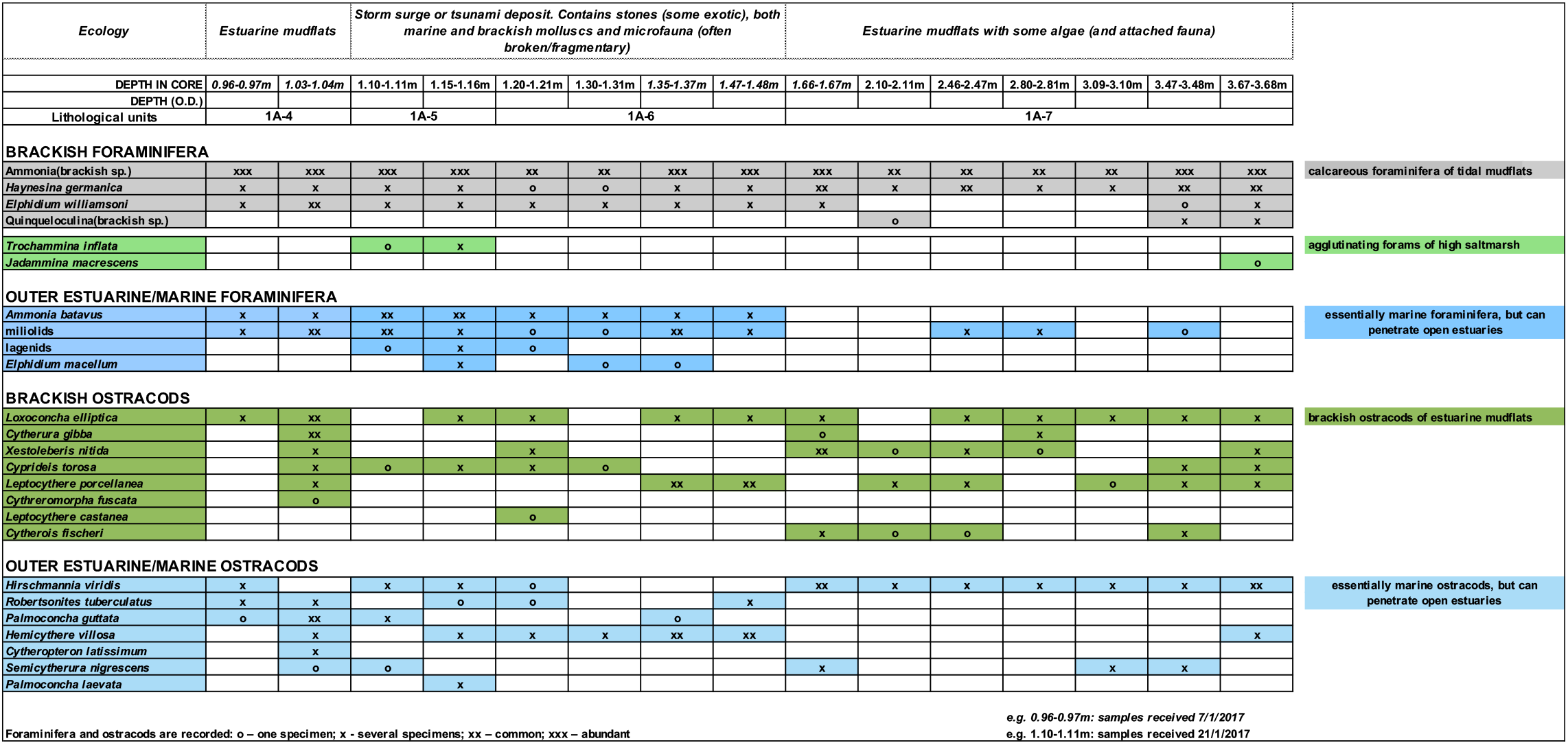
Foraminifera and ostracod abundance profiles of core ELF001A.

#### S4.2 Pollen analysis

Subsamples of 1cm^3^ were extracted from the core at 0.05m intervals and prepared for pollen analysis using standard methodologies, including HF treatment and Acetylation. *Lycopodium* spores were added to permit the calculation of pollen concentrations. Pollen counting was carried out on a Leica DM100 at a magnification of x400. All pollen nomenclature follows Moore *et al.* (1991)^52^ with the amendments proposed by Bennett *et al.* (1994)^53^. At least 150 pollen grains were counted per sample.

The results are presented as a pollen diagram produced using TILIA and TILIA*GRAPH^54^ (Figure 4, Figure S4.1A). Microscopic charcoal fragments were counted and are expressed as a percentage of total land pollen. The diagram has not been divided into biostratigraphic assemblage zones, but the position of the *Tsunami* deposit between 1.03-1.55m is indicated, dividing the sequence into pre- and post-*Storegga*; no pollen was preserved in this unit (Unit ELF001A-6). The diagram is dominated by relatively few, predominantly arboreal taxa: total tree and shrub percentages are generally above 90% total land pollen (TLP) with herbs accounting for a maximum of 20%. This implies the presence of dense woodland in the pollen source area. *Corylus avellana*-type (likely to be hazel, rather than *Myrica gale* in this situation), is dominant throughout (c. 60% TLP). Other trees which are consistently recorded but at lower percentages, are *Quercus* (oak) and *Pinus sylvestris* (Scots’ pine) (both c. 10-20%), with lower values for *Ulmus* (elm; up to 5%), *Alnus glutinosa* (black alder; c. 5%) and *Betula* (birch, max 9%). Other trees/shrubs recorded sporadically at low percentages are *Tilia* (lime), *Salix* (willow), *Fraxinus* (ash) *Hedera helix* (ivy) and *Ilex aquifolium* (holly).

Herbaceous taxa account for a relatively low proportion throughout, but with Poaceae (wild grasses) consistently present (max 15%). Another herb recorded in almost every sample (max 5%) is *Silene dioica*-type (red campion), whilst Cyperaceae (sedges), Chenopodiaceae (Fat Hen family), *Artemisia*-type (mugwort). Ranunculaceae (buttercups), *Filipendula* (meadowsweet) and a few other herbs make occasional appearances including *Sedum* (stonecrop). Spores including Pteropsida (monolete) indet. (ferns), *Polypodium vulgare* (common polypody), *Sphagnum* (bogmoss) and *Pteridium aquilinum* (bracken) are present throughout, with the former best represented (max 8% TLP+spores). Proportions of microscopic charcoal are rather variable, but seem to be higher in the uppermost three samples of the sequence. The spectra are remarkably consistent, other than for a spike in *Quercus* at 2.65m, associated with a reduction in *Corylus*. It is difficult to assess what processes this temporary expansion of oak relates to as there are no other pronounced changes at this level.

Overall, the sequence indicates a landscape of deciduous woodland, in which hazel was dominant with oak, pine, elm and birch as subordinate components, and alder and willow on damper soils. The impression of a shady, closed woodland is reinforced by the presence of ivy and common polypody, often found as an epiphyte on oak trees. The consistent record of grass throughout may reflect the presence of open areas within the woodland, but more probably reflects the presence of wetland grasses such as *Phragmites* (reeds) growing in the lagoonal environment. The range of herbs also indicate communities typical of damp soils (buttercups, sedges, meadowsweet), perhaps growing on the ecotonal areas between the lagoon and the dryland. In particular, the consistent presence of *Silene dioica*-type is notable; the probable species represented is *Silene dioica* which typically grows in partially shaded habitats, also indicated by the record of fern spores. The Chenopodiaceae includes many herbs, but in this context is most likely to indicate plants of this family that grow on salt marshes and other saline soils; the rare records of stonecrop perhaps also reflect drier, sandy soils typical of the coast. The presence of *Pteridium* might also imply better drained soils in the wider landscape where bracken would be found. In general there is no evidence for any form of disturbance to the environment for the duration of the record.

The most striking aspect of the sequence is the relative lack of fluctuation throughout, the curves of all the taxa are remarkably stable. Two comments are pertinent to how this relative homogeneity might be interpreted. Firstly, the data indicate that the vegetation within the pollen source area was broadly stable across the period of time represented by the diagram with no palynologically identifiable changes. Secondly, it is possible that this apparent stability of the environment through time, is related to the nature of the pollen source area for the silt dominated deposits that constitute the sequence. It is likely the pollen derived from a relatively large spatial area, including the dryland landscape adjacent to the lagoon but also terrestrial locations further upstream. In other words, the pollen record is resolving an area of landscape of potentially tens of square kilometres. Moreover, there is likely to have been a degree of mixing and reworking of the pollen within the water column, so interpretation must of necessity be tentative in terms of the extent or character of inferred vegetation dynamics throughout the sequence.

There are no pronounced changes in the spectra immediately above the hypothesised *Tsunami* layer, which might be taken to imply that this event had no identifiable impact on the local vegetation, with a predominantly wooded landscape both pre and post-*Tsunami*. However, this interpretation must be tempered by the previous comments concerning the potential taphonomic complexity of the pollen record. However, there is evidence of potential changing woodland dynamics above this unit, towards the top of the diagram, between 1.03-0.84m. Total tree pollen percentages increase as a result of rising values for *Pinus* alongside reductions in *Corylus*. Again, this is difficult to interpret but may indicate an expansion in Scots pine, prior to the establishment of marine conditions at this location. Interestingly, there are also increased representation of microscopic charcoal across the same levels. It would be tempting to interpret these changes as potential evidence for increased dryness in the period before the final marine incursion at this location. Otherwise, there is no palynological evidence for woodland recession that would be expected to result from rising water tables in advance of rising relative sea levels.

**Figure S4.1.**
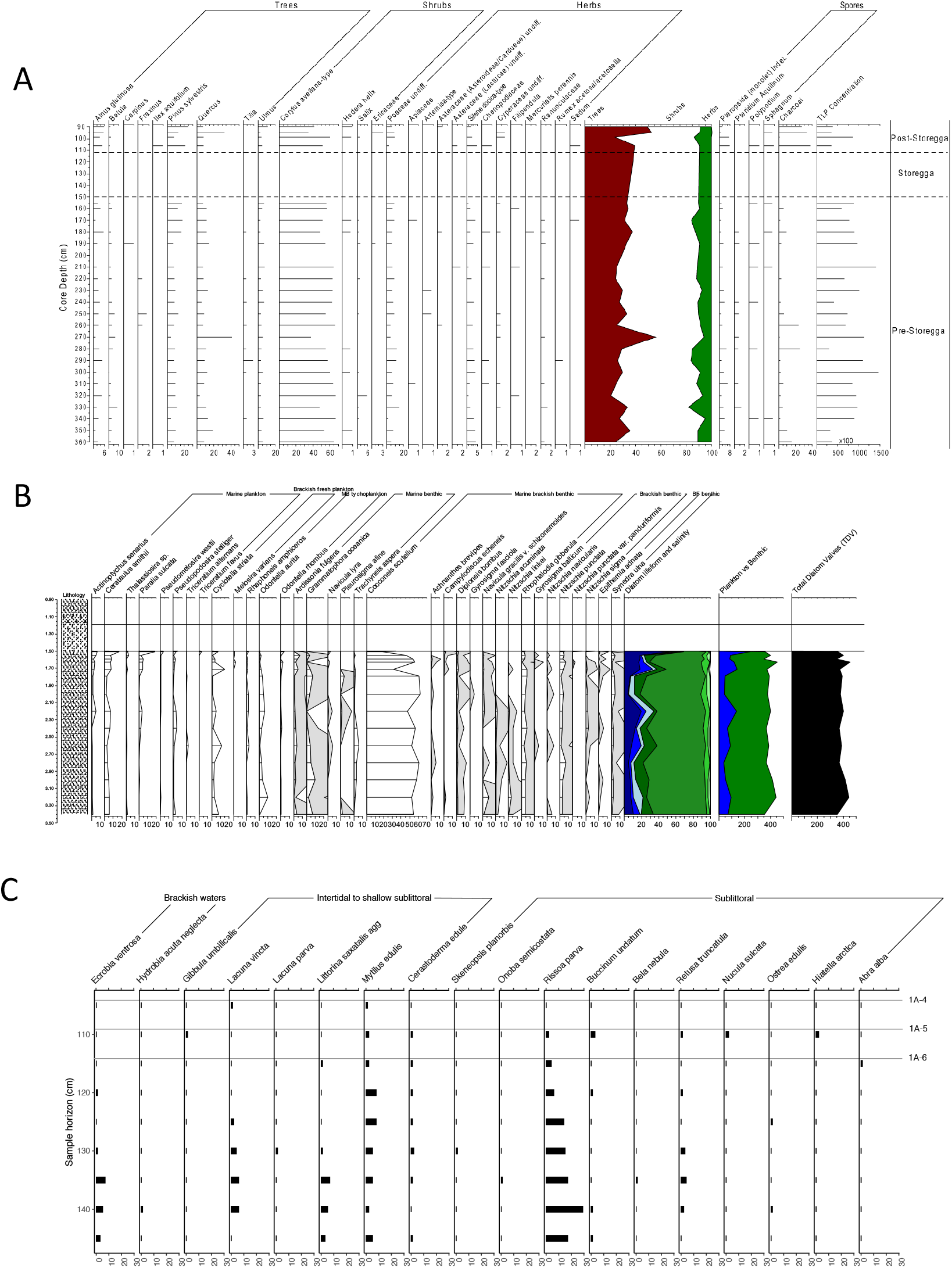
ELF001A palaeoenvironmental proxy count profiles. All measurements taken from lithological Unit ELF001A-4 and below. A. pollen counts B. diatom counts C. mollusc counts.

#### S4.3 Diatom analysis

A selection of 23 spot samples were prepared for initial diatom assessment from the sedimentary sequence of core ELF001A. Diatom preparation followed the methodology of Plater et al. (2000)^55^, with additional pretreatment using sodium hexametaphosphate, to assist in minerogenic deflocculation. Samples were sieved using a 10μm mesh to remove fine minerogenic sediments. The residue was transferred to a plastic vial, from which a slide was prepared, using Naphrax as the slide mountant, for subsequent assessment.

For samples in which diatoms were encountered in sufficient abundance during the initial assessment, a minimum of 300 diatoms were identified for each sample depth. If preservation was found to be poor, a complete slide was traversed in an attempt to extract the diatom data available from the sample under assessment. Poor preservation was experienced in the majority of samples from within and above the event stratum. Diatom species were identified with reference to van der Werff and Huls (1958-74)^56^, Hendy (1964)^57^ and Krammer & Lange-Bertalot (1986-1991)^58^. Ecological classifications for the observed taxa were then achieved with reference to Vos and deWolf (1988; 1993)^59,60^, Van Dam et al., (1994)^61^ and Denys (1991-92; 1994)^62,63^.

The overall diatom signal from within the sediments underlying the event stratum can be interpreted as indicating coastal conditions prevailing throughout its depositional history (Figure 4, Figure S4.1B). The dominance of marine to brackish benthic taxa, with a particular presence of taxa often associated with plants and muddy substrates (epiphytic and epipelic taxa respectively), would infer deposition took place within the intertidal zone. The absence of aerophilous taxa and epipsammic taxa is also noted, whilst planktonic/tychoplanktonic taxa rarely contribute more than 20-30% Total Diatom Valves (TDV) to the floral assemblages. The diatom assemblages are also found to be consistent throughout the sedimentary unit, which suggests similar conditions prevailed throughout the deposition of the sedimentary unit.

The dominant taxa, *Cocconeis scutellum,* is a ‘northern’ epiphytic species often encountered in the littoral zone of the North Sea and Arctic oceans^64^, and is affiliated with taxa such as *Zostera marina* or seagrass (Main & McIntyre, 1974; cited in Werner, 1977)^65^ as well as green algae such as Cladophora sp.^66^. Studies by Tanaka (1986)^67^ have also shown that *C. scutellum* is commonly associated with seaweed (including *Sargassum horneri*, *S. patens*, *S. piluriferum*) and *Undaria* (kelp). When combined with the relative dominance of other epiphytic taxa throughout the sedimentary profile, we can first infer that deposition relatively high on the tidal frame. The low but persistent presence of marine planktonic and tychoplanktonic species indicates tidal inundations occurred, but were somewhat restricted, during the development of the deposits underlying the event stratum.

When comparing such floral assemblages encountered beneath the event stratum to the ecological groupings stipulated by Vos and deWolf (1993)^60^, deposition within a setting which experiences a large tidal range is discounted. This is due to the absence of any aerophilous and epipsammic taxa, the relative dominance of marine-brackish epiphytic taxa and the relatively limited influence of planktonic and tychoplanktonic taxa. A microtidal palaeoenvironment such as a tidal lagoon or small tidal inlet is interpreted. All samples contain diatoms typical of such a depositional setting, to infer the environment remained stable throughout the deposition of the finely laminated silt and clays that underlie the event stratum.

#### S4.4 Mollusca analysis

Nomenclature followed WoRMS (WORMS EDITORIAL TEAM 2018)^68^. Identifications were carried out using a reference collection. Ecological information is derived from Graham (1971)^69^ and Allcock et al. (2017)^70^. Minimum number of individuals (MNI) for gastropods was determined by counting all non-repeating elements of that species within a sample and using the largest number as the MNI. In the case of bivalves, only shell hinge fragments or intact valves were counted. Minimum numbers of left and right valves are presented separately. The highest of these two numbers is used as the MNI.

Preservation was largely good in all samples, although *Mytilus* shells and other bivalves were almost invariably broken (Figure S4.1C). Numbers of shells were generally low, although shells are more frequent within Unit ELF001A-6. Numbers decline through time. At the bottom of the sequence, in Unit ELF001A-6, there is an ecologically mixed assemblage. There is a brackish water fauna present in this unit as well, represented by moderate numbers of *Ecrobia ventrosa,* and a single *Hydrobia acuta neglecta.* These are snails associated with relatively low salinities, in sheltered locations such as estuaries and lagoons, away from high energy tidal influence. The same samples also contain low numbers of fruits of *Potamogeton spp.* (pondweed), which is found in fresh to brackish water settings. The unit is dominated, however, by taxa from a lower shore or sublittoral environment, especially *Rissoa parva.* This is a common snail under stones and on weeds from mid tidal level down to 15m depth on rocky shores. Other molluscs present in this unit include *Retusa obtusata,* a predatory gastropod which was most likely preying on *Rissoa;* and the common mussel *Mytilus edulis,* which is usually found intertidally on rocky coasts. The mussel shells are all broken, which may suggest compression from overlying sediment or wave transport, however they are not especially rounded, which suggests they were not subjected to much wave rolling. Rather than rocky shores, common cockle, *Cerastoderma edule,* and European oyster, *Ostrea edulis,* are found in muddy and sandy environments. The intertidal to lower shore assemblage continues to dominate in Unit ELF001 A-5, however the brackish water fauna is now absent. There is lower species diversity in this unit, however there is somewhat more equitability, with *Rissoa* less dominant. In unit ELF001A-4, numbers are very low, containing just a single shell each of *Lacuna vincta* and *Mytilus edulis,* which appear to reflect an intertidal setting on a high-energy coast. Overall, the samples appear to suggest inundation of a previously low energy tidal-dominated estuarine setting and establishment of a much higher energy wave-dominated environment. The transition is not clear however, and ecological signals remain mixed throughout Unit ELF001A-6, indeed the brackish water fauna peaks at 1.35-1.40m depth. A likely scenario is that this deposit represents a conflation of material eroded and reworked from markedly different locales (rocky, wave-dominated; and muddy, tidally-dominated). The complete absence of abrasion on the shells suggests that they were not subject to usual wave transport. Bivalve shells with angular breaks that have not been wave-rounded may be features of tsunami deposits^71^, however in these samples there are no articulated bivalves, which Donato et al. also found in a recent tsunami deposit from Oman

#### S4.5 SedaDNA analysis

##### DNA extraction

All DNA handling stages prior to PCR took place in a dedicated aDNA facility at the University of Warwick following standard protocols for processing ancient DNA^72^. Sealed sediment cores were refrigerated at 4°C immediately after retrieval and were held at a constant 4°C until sampling. Cores were split under strict aDNA lab conditions and under red-light to preserve samples for OSL analysis. All samples for aDNA work were taken inside a category two biosafety cabinet using sterile equipment. The cut surface of the core was removed and ∼20 g of sediment retrieved, ensuring that no sediment from the outer 1 cm of the core was collected, as this may have been disturbed during the coring process. For DNA extraction, library preparation, sequencing, and downstream analysis, each sample was processed in duplicate. For each duplicate, 2 g (±0.05 g) of sediment was taken from each sample. Subsamples were processed in batches of up to seven plus one negative control (reagents only). The subsamples were mixed with 5 ml CTAB buffer (2% w/v CTAB, 1% w/v PVP, 0.1 M Tris pH 8.0, 20 mM EDTA, 1.4 M NaCl) and incubated at 37°C with agitation for 7 days. After incubation, the subsamples were centrifuged at 20,000 xg for 10 minutes. The supernatant was moved to a new 50 ml tube and manually shaken with 4 ml chloroform:isoamyl alcohol (24:1) for 5 minutes. The resulting mixture was centrifuged at 20,000 xg for 5 minutes. The aqueous phase was combined with 20 ml Buffer AW1 (Qiagen) and incubated at room temperature for 1 hour. This was then applied to silica-based spin columns using a vacuum manifold. The columns were then washed, first with 500 μl Buffer AW2 (Qiagen), and then 300 μl acetone, both followed by centrifugation at 6,000 xg for 1 minute. The columns were then removed from their collection tubes and air dried for 5 minutes. Finally, DNA was eluted in 65 or 75 μl Buffer EB (Qiagen). They were incubated at 37°C for 10 minutes and centrifuged at 15,000 xg for 2 minutes. The eluted DNA was quantified using a high-sensitivity Qubit assay (Invitrogen).

##### Sequence generation

The library protocol is based on Meyer and Kircher (2010)^73^ with the following modifications from Kircher et al. (2012)^74^: 0.1 μl of adapter mix during adapter ligation instead of 1 μl; spin column purification (MinElute PCR purification kit, Qiagen) instead of SPRI; purification step after adapter fill-in replaced with heat inactivation for 20 minutes at 80 °C; Double indexing; no fragmentation step, as ancient DNA is expected to already be shorter than 400 bp; blunt-end repair reaction volume of 40 μl; T4 DNA ligase added to individual sample tubes instead of the master mix during adapter ligation; Platinum Pfx was used indexing PCR for most samples, but since this was discontinued in 2018, Platinum SuperFi was used for some samples; there were 16 PCR cycles for most samples, but where Platinum SuperFi was used 18 PCR cycles were applied.

Libraries were visualised on a 2% agarose gel. They were then cleaned using 45 μl SPRI beads and eluted in 20 μl TET buffer^75^. The cleaned libraries were quantified using a Qubit assay (Invitrogen) and a fragment size profile produced using a Bioanalyzer (Agilent). Libraries were normalised to 4nM and pooled prior to sequencing on the Illumina NextSeq platform using the high-output, v2, 150-cycle kit (75×75 paired end), Table S4.2. Sequence data were deposited in the European Molecular Biology Laboratory European Bioinformatics Institute (project code PRJEB33717).

##### Bioinformatics

Raw BCL files were converted to FASTQ and demultiplexed using Illumina’s bcl2fastq software (version v2.20.0.422), using the --no-lane-splitting and --ignore-missing-bcl options. Adapters were removed and paired end reads were collapsed using AdapterRemoval (version 2.2.2)^76^, specifying a minimum length of 30 and a minimum quality of 30. FastQC (version 0.11.6)^77^ was used to visually assess the success of adapter and quality trimming. FASTQ reads were converted into FASTA format using the following example shell command: In.fastq | awk ‘NR%4 !=0’ | awk ‘NR%3 !=0’ | sed ‘s/@/>/g’ > out.fasta. Finally, duplicates were removed using the fastx_collapser command from the FASTX-toolkit (version 0.0.13)^78^.

An initial metagenomic BLASTn search (version 2.6.0)^79^ was undertaken using the tab output (specified using -outfmt “6 std staxids”). This allows a large volume of data to be processed with a far smaller data footprint than the full BLAST output format. This was then converted to RMA format using the MEGAN5 command line (version 5.11.3)^80^, enabling the visualisation of the preliminary data. The patchiness of DNA sequence databases and the overrepresentation of model organisms leads to unreliable assignation of sequences. Reads were therefore stringently filtered using the Phylogenetic Intersection Analysis (PIA)^81^. FASTA sequence reads with preliminary assignation to taxa of interest (in this case Viridiplantae, Chordata with primate reads excluded, Arthropoda, and a random subset of 10,000 bacterial reads) were extracted from the RMA files using MEGAN5 command line tools (version 5.11.3)^80^, These reads were subjected to a second round of BLASTn (version 2.6.0)^79^, this time to generate the full BLAST format as an output. These were then used as an input for Phylogenetic Intersection Analysis (Smith et al., 2015; default settings) ^81^ in order to retrieve stringent assignations. Taxa with >2% of the assigned reads also assigned to that taxon in the negative controls for that sequencing run were removed. Any taxa that remained after the stringent filtering that were not native to Europe were discarded, accounting for about 3% of the data.

##### Authentication: DNA damage analysis

Two approaches were taken to establish whether damage patterns characteristic of ancient DNA were present to authenticate the sedaDNA. Current damage authentication methodology is predicated on the reconstruction of ancient genomes^16^, rather than metagenomic assemblages present in shotgun data where the coverage of any one genome may be too low for a significant signal. Fortunately, a number of tree species from the Unit ELF001A-6 were well enough represented to use this approach, *Quercus*, *Betula* and *Corylus*. Paired end collapsed reads were mapped to *Betula*, *Corylus*, and *Quercus* genomes (accessions: GCA_900184695.1, C.avellana_Jefferson OSU 703.007, and GCA_900291515.1 respectively) using BWA-ALN (version 0.7.12-r1039)^82^ specifying - n as 0.01 and -l as 1000. BAM files were generated using bwa samse (version 0.7.12-r1039)^76xx^, and samtools view with the -Sb flag (version 1.7) ^83^. Read group tags were added using picard AddOrReplaceReadGroups (version 2.18.7) (http://broadinstitute.github.io/picard/), then duplicates were marked and removed using picard MarkDuplicates. Realignment around indels was undertaken using the GATK tools RealignerTargetCreator and IndelRealigner (GATK version v3.8-1-0-gf15c1c3ef) ^84^. All intermediary sorting and indexing stages were undertaken using the samtools sort and index tools (version 1.7) ^83^. MapDamage (version 2.0.6)^16^ was then used to assess the extent of DNA damage patterns using the –merge-reference-sequences option. Fragment misincorporation plots can be seen in Figure S4.2. The fragmentation parameter λ was calculated for these genome mapped data sets using the methodology of Kistler *et al.* 2017^15^, Figure S4.3.

A second novel methodology was also applied which does not require mapping DNA to a genome to estimate misincorporation parameters, to allow the assessment of a taxonomically mixed assemblage. The metagenomic damage analysis tools allows for the assessment of post-mortem deamination patterns on a metagenomic scale (i.e. whole sequence sample files), instead of an individual reference genome comparison with individual hits, used with tools such as MapDamage^16^. Metagenomic damage analysis is based on a three-step bioinformatic process, and examines the first 5’ 19 base pair positions. All sequences were subjected to metagenomic BLASTn analysis^79^ with the ‘qlen’ option, using the full NCBI nt database. The ‘qlen’ option creates a standard output with the known available positions and the full sequence length for each hit. Using the BLAST outputs, a combined fasta file of the hit IDs and associated reference genome sequence was then created using Efetch (part of the Enterez direct tool) within the E-Utilities package, which provides access to the NCBI’s suite of interconnected databases. This process was piped into PERL script ‘Efetch.pl’, which is a four-step process. The script firstly opens the BLAST output and, on a conditional argument, builds in a directional variable which sorts the base-pair start and end position depending on whether the read is in non-reversed or reverse complemented alignment. Once the start and end coordinate of each hit is sorted, the script using the efetch command, connects to the NCBI database and copies the associated organism information and positional read from the reference genome. If in the reverse complement, the next step is to reverse it to the original sequence alignment. Finally, the hit ID, organism information and the matched section of the reference genome are piped into individual FASTA files for each of the original BLAST ID hits. Once each individual FASTA file is created, using PERL script ‘newBlastParse.pl’, the sequence from each hit from the original BLAST output was then appended to the associated individual FASTA file, creating a two record FASTA. This process finds the query ID, subject ID and the sequence from the original BLAST output, and prints into the associated FASTA file created in the previous step. Once a fully populated FASTA file for each individual BLAST hit was created, the Efetch record and the BLAST ID were realigned using the Needleman-Wunsch algorithm^85^, using the ‘aln.pl’ which records the alignment of each base pair. Once aligned, each hit was assessed for positional mismatches using PERL script ‘mismatches.pl’, which examines each base position and counts the prevalence of any C > T misincorporations. The positional mismatches were visualised in RStudio (V1.1.456).

The constant 4°C environment of the sea floor leads to an expectation of damage which may be as low as 2.5% of terminal overhang cytosines deaminated in the age ranges explored in this study^15^, which may be below levels of detectability as has been observed in previous studies^81,86^. Furthermore, the high ionic environment of marine conditions is expected to reduce deamination rates by reducing the rate of hydrolytic attack^87,88^, as has been observed for marine environments^89,90^. Here we observed deamination rates in the range of 7-15% which is in line with these expectations of low damage levels for the sediments of Unit ELF001A-6, dated to 8.14 ± 0.29 Kyrs. BP (Figure S4.2).

The metadamage analysis across the sedaDNA set confirms this low level of damage signal, agreeing closely with the mapdamage assessment indicating damage levels are reflected across taxa (Figure S4.4). We applied the metadamage analysis to sequence data both before and after the PIA filtration step. The mismatch base line in the post PIA analysis data is lower than the pre-filtered data, indicating a lower level of phylogenetic background noise as would be expected as less accurate phylogenetic assignations are rejected. In this way the metadamage analysis validates the PIA analysis.

Fragmentation profiles indicate a high variance within horizons indicating poor correlation with age as has previously been observed^15^ (Figure S4.3).

##### Biogenomic mass

Shotgun data have the potentially useful property of representing the biomass of organisms present in terms of cell counts, if genome size is taken into account. We used C values of the Kew Angiosperm database (http://www.kew.org/cvalues/) to estimate representative genome sizes for floral taxonomic units, while we used the Animal Genome Database (http://www.genomesize.com) for faunal estimates. We calculate a biogenomic mass value by dividing the number of sequence read counts observed for a taxon by the genome size to give a counts per gigabase value. This value should give an estimate that is directly proportional to the number of cells left behind by organisms, assuming minimal effects of clonal bias in library preparation, Figure 4. Both raw count values and associated biogenomic mass values are represented for floral (Figure S4.5) and faunal (Figure S4.6) for core ELF001A, and values for floral profiles are shown for cores ELF003, ELF0031A, ELF0039 and ELF059A (Figure S4.7).

##### Authentication: stratification analysis

Studies of sedaDNA need to establish the stratigraphic integrity of ancient DNA recovered, and whether there has been post deposition movement of DNA up and down the sediment column as has been observed in past studies^91^. To date, no clear methodologies have been established for best practice to check for DNA movement. Here we applied a statistical approach to calculate the probability that taxon counts between horizons could have been drawn from the same statistical distribution, indicating homogeneity of DNA titre across horizons and therefore complete diffusion. While complete diffusion represents the extreme of DNA movement, the probability never the less provides a metric of the abruptness of change allowing an evaluation of the likelihood horizon pairs are part of the same diffused population. The methodology is equally applicable to other biological proxy data such as pollen and diatoms, and so was applied to the data sources in this study. Taxon counts are assumed to follow a Binomial distribution where the total count number represents the trial number, and the taxa count is the number of observed successful outcomes. We then applied Beta distributions to explore the underlying probability of the Binomial distributions where parameters a and b were derived from the number of counts of a taxon and the total number of counts minus the taxon counts respectively. The probability that two sets of taxon counts were derived from the same underlying distribution (*p value*) was inferred from the area of overlap between the two derived Beta distribution probability density functions (Figure 4, Figure S4.8).

We further quantified the extent of change in taxon count number between by applying an index of change (Figure 4, Figure S4.5) outlined in equations 1 and 2:

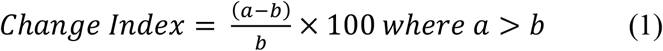

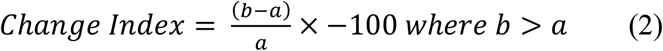

Where a and b are the maximum likelihood estimators of the probability of a count being assigned to a particular taxon as derived from the Beta distribution of the overlying and underlying horizons respectively.

Highly significant differences were observed between horizons, indicating a lack of movement of DNA in the sediment column. In the case of Unit ELF001A-6 and the underlying Unit ELF001A-7, *p* values for woody taxa such as Fagales (9.9 x 10^−51^), Quercus (5.2 x 10^−5^), Betula (5.96 x 10^−25^), Saliliceae (7.63 x 10^−57^) and the Amygdaloideae (1.17 x 10^−257^) are highly convincing of an abrupt change and therefore lack of DNA movement, Figure 4. Note in the case of ELF003 radio carbon dates indicate an inversion in which the underlying tsunami associated unit is younger than the overlying units, hence the significant fall in tree taxa comparing these two units (Figure S4.8) should be interpreted as a significant influx of woody taxa associated with the tsunami.

**Figure S4.2.**
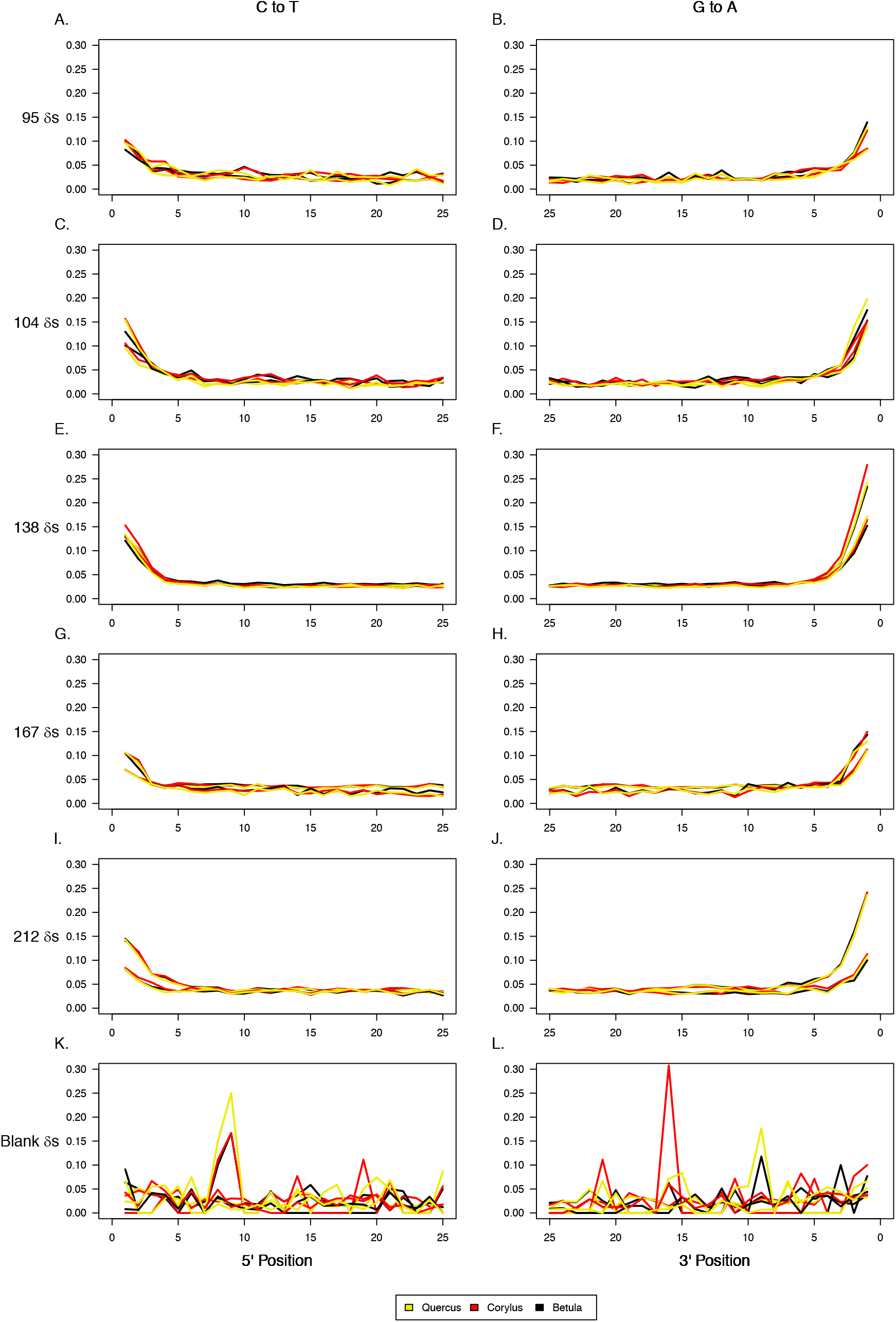
C to T mismatch distributions of sedaDNA mapped to genomes. *Quercus*, *Corylus* and *Betula* genomes used, calculated in mapDamage 2.0^16^

**Figure S4.3.**
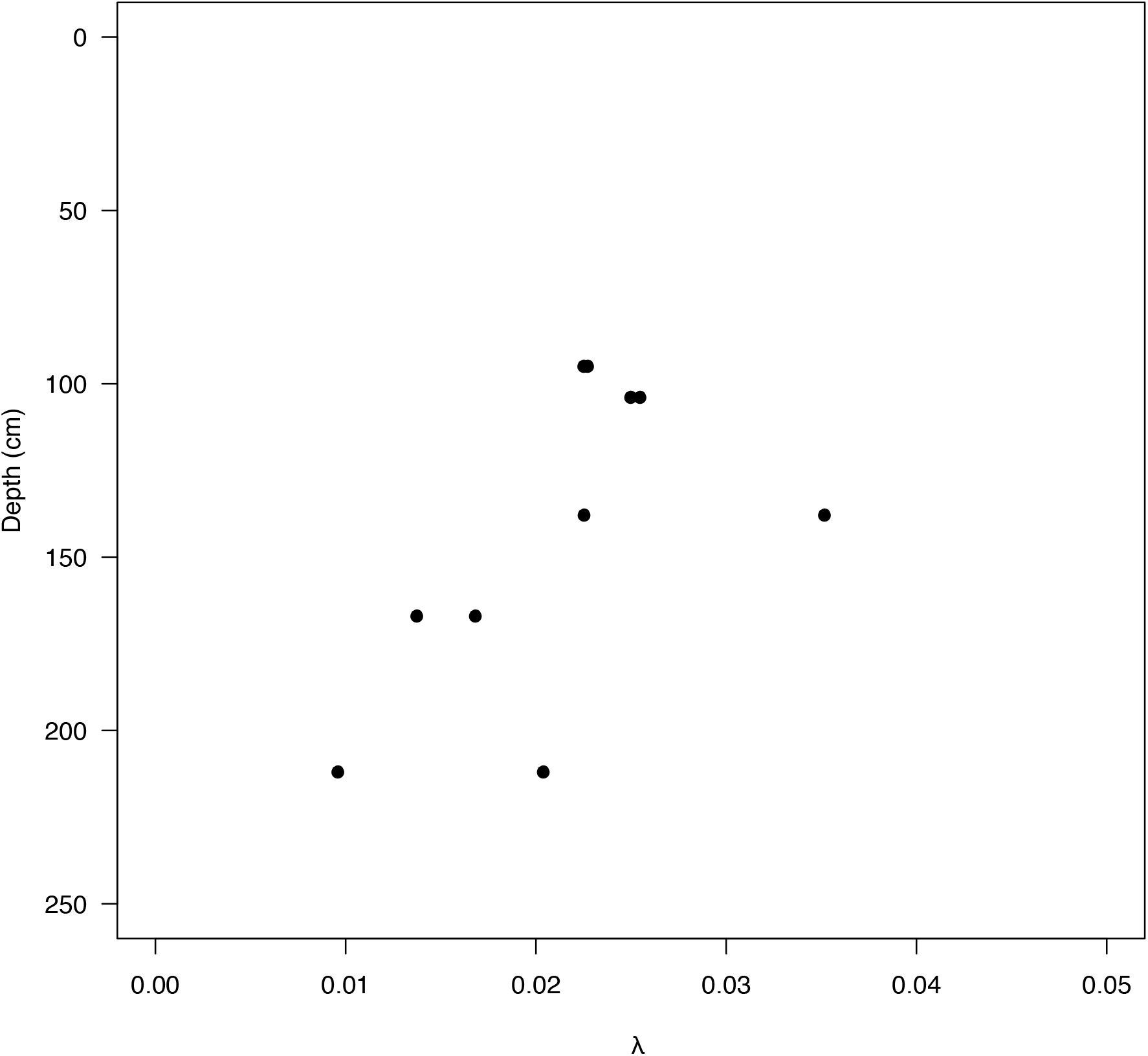
DNA fragmentation pattern derived λ statistics for core ELF001A.

**Figure S4.4.**
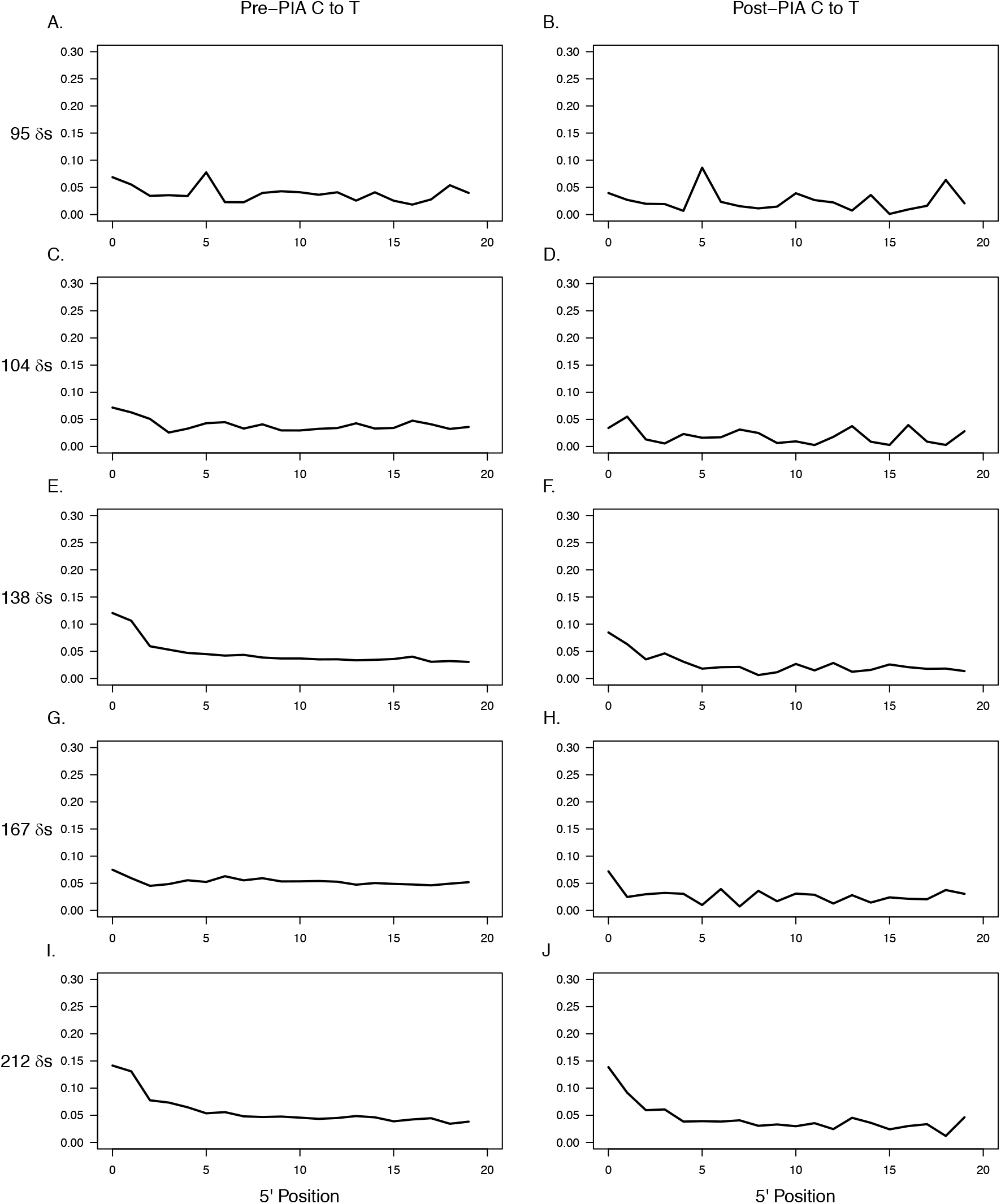
Metadamage analysis of C to T transitions across all pre and post PIA filtered sedaDNA

**Figure S4.5.**
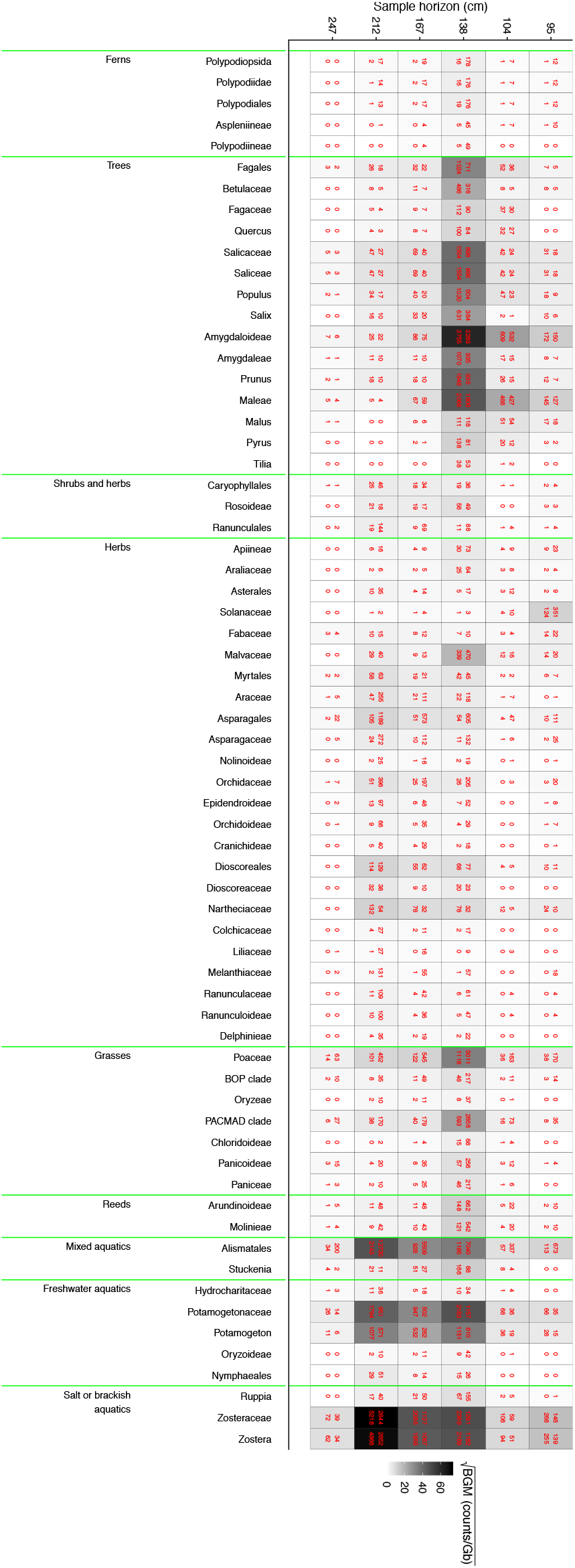
ELF001A floral sedaDNA profile. Numbers above are absolute counts of reads after PIA filtering, numbers below are biogenomic mass (BGM), counts/Gb. Shading represents corresponding BGM values scaled by square root.

**Figure S4.6.**
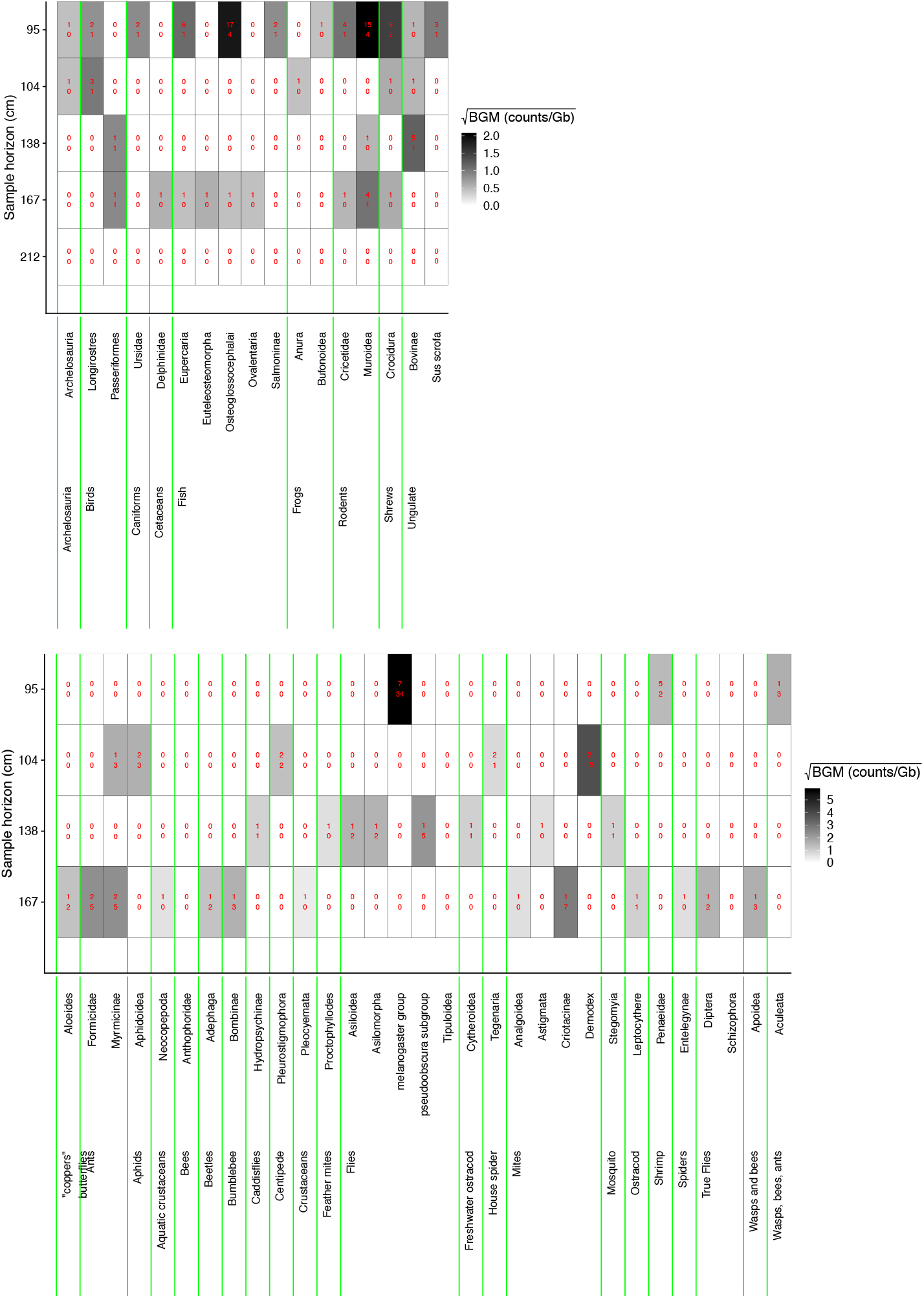
ELF001A faunal sedaDNA profile. Numbers and shading as Figure S4.5. Upper panel: vertebrates, lower panel: invertebrates.

**Figure S4.7.**
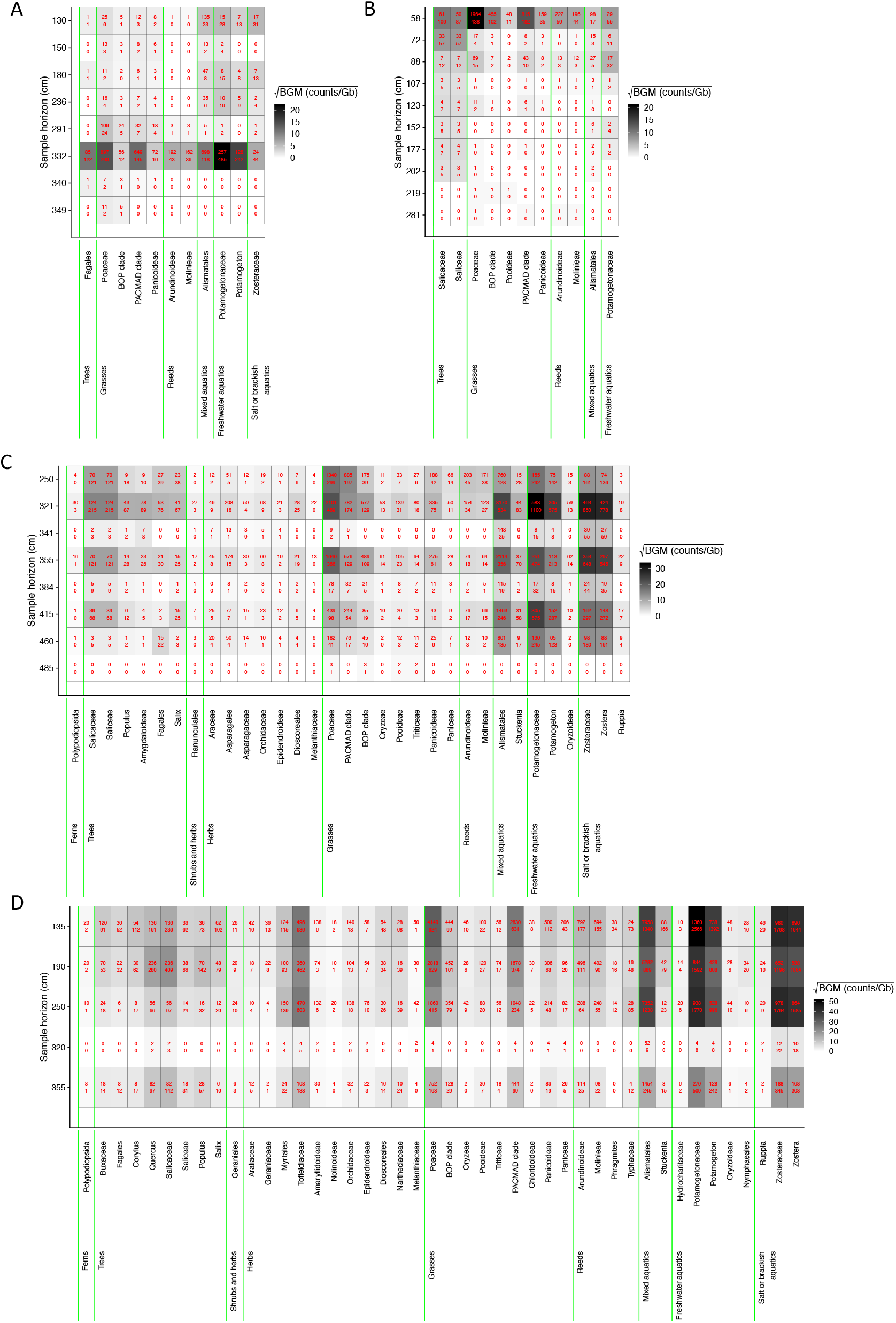
Floral sedaDNA profile of other tsunami candidate cores. Numbers and shading as Figure S4.5. A. ELF003, B. ELF0031A, C. ELF0039, D. ELF0059A.

**Figure S4.8.**
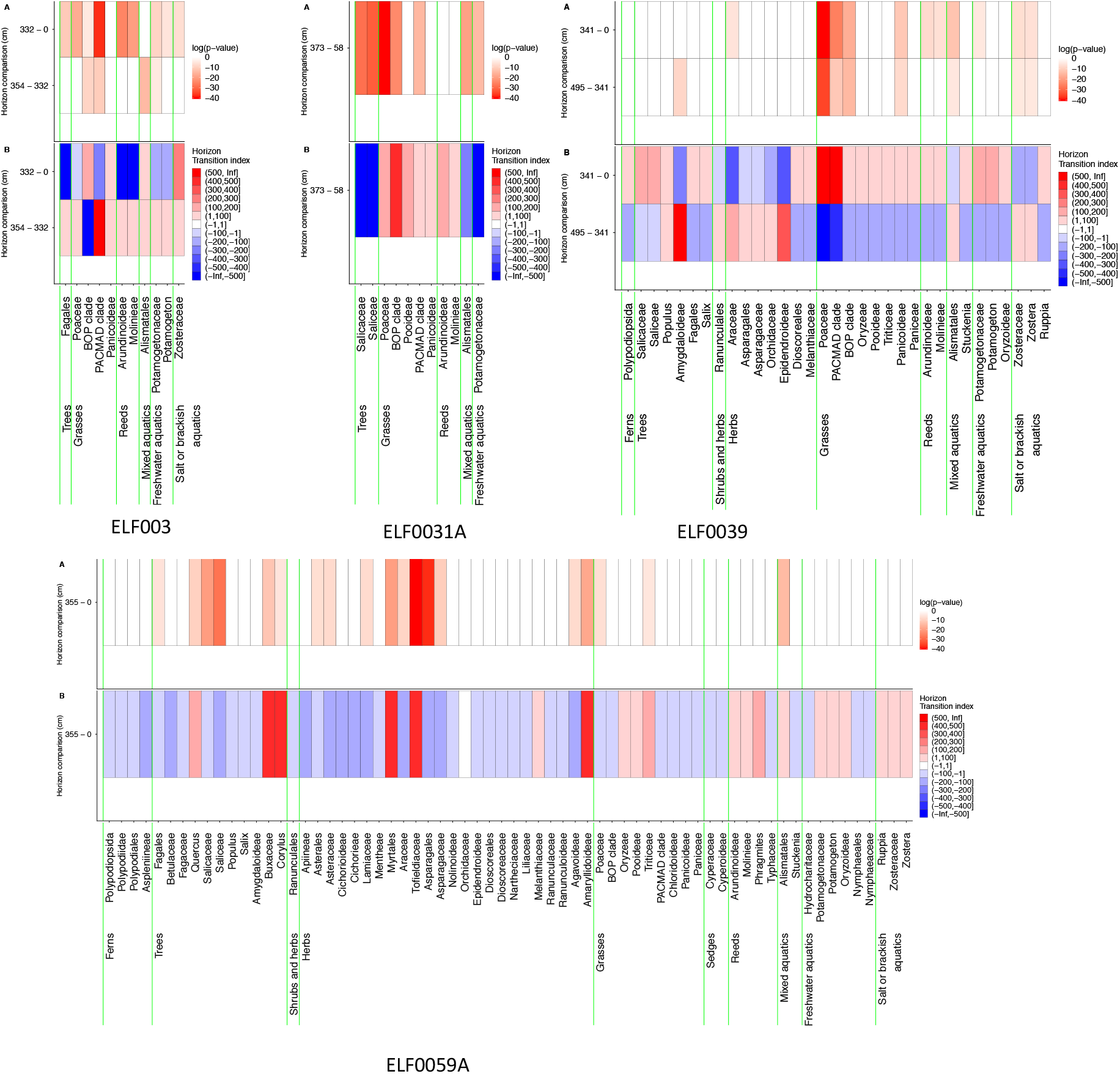
sedaDNA change between putative tsunami and adjacent strata. Assessment of taxon change between sample horizons of taxa with abundances > 50 in other tsunami candidate cores identified by seismic survey. Below: Index of change between horizons based on changes in maximum likelihood estimators of the probability taxon being selected from each horizon. Blue indicates a decrease in probability moving up the core, red an increase. Above: Probability of observed taxa counts between pairs of horizons being drawn from the same distribution. A. ELF003, B. ELF0031A, C. ELF0039, D. ELF0059A.

**Table S4.2.**
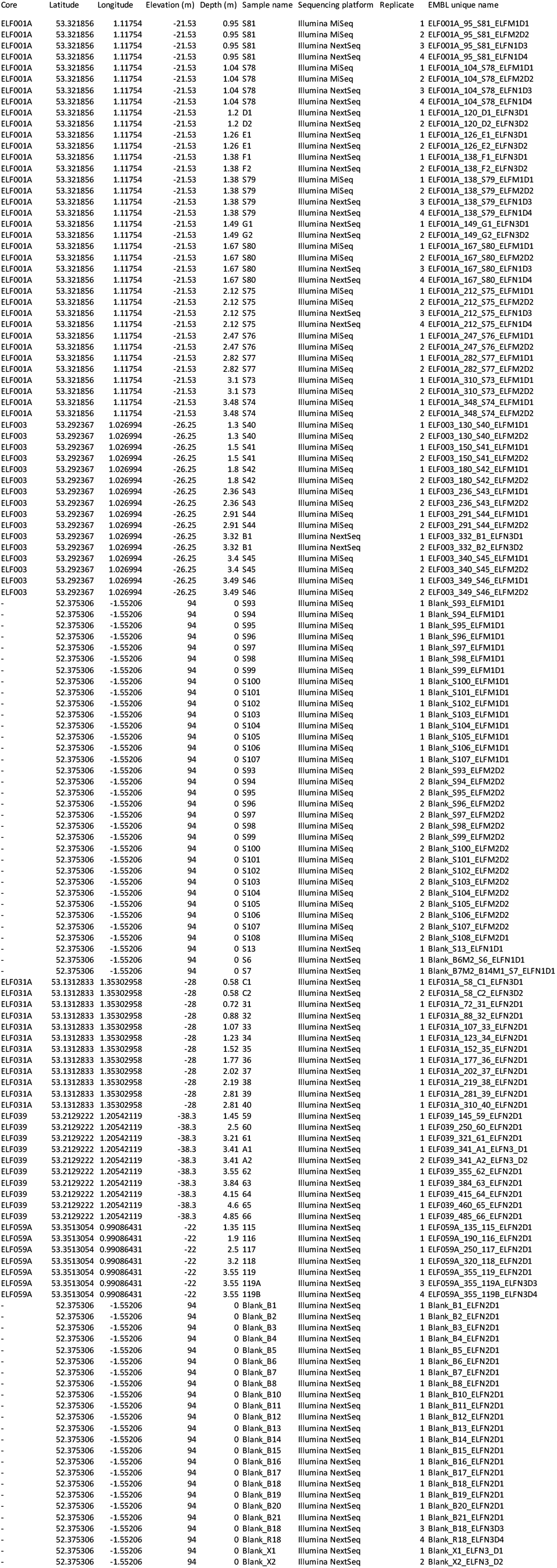
SedaDNA sample and read details.

